# Activation of XBP1s attenuates disease severity in models of proteotoxic Charcot-Marie-Tooth type 1B

**DOI:** 10.1101/2024.01.31.577760

**Authors:** Thierry Touvier, Francesca A. Veneri, Anke Claessens, Cinzia Ferri, Rosa Mastrangelo, Noémie Sorgiati, Francesca Bianchi, Serena Valenzano, Ubaldo Del Carro, Cristina Rivellini, Phu Duong, Michael E. Shy, Jeffery W. Kelly, John Svaren, R. Luke Wiseman, Maurizio D’Antonio

**Author notes:** Correspondence to: Maurizio D’Antonio, Division of Genetics and Cell Biology, IRCCS Ospedale San Raffaele, Via Olgettina 58, 20132, Milan Italy.

## Abstract

Mutations in myelin protein zero (MPZ) are generally associated with Charcot-Marie-Tooth type 1B (CMT1B) disease, one of the most common forms of demyelinating neuropathy. Pathogenesis of some MPZ mutants, such as S63del and R98C, involves the misfolding and retention of MPZ in the endoplasmic reticulum (ER) of myelinating Schwann cells. To cope with proteotoxic ER-stress, Schwann cells mount an unfolded protein response (UPR) characterized by activation of the PERK, ATF6 and IRE1α/XBP1 pathways. Previous results showed that targeting the PERK UPR pathway mitigates neuropathy in mouse models of CMT1B; however, the contributions of other UPR pathways in disease pathogenesis remains poorly understood. Here, we probe the importance of the IRE1α/XBP1 signalling during normal myelination and in CMT1B. In response to ER stress, IRE1α is activated to stimulate the non-canonical splicing of *Xbp1* mRNA to generate spliced *Xbp1* (*Xbp1s*). This results in the increased expression of the adaptive transcription factor XBP1s, which regulates the expression of genes involved in diverse pathways including ER proteostasis. We generated mouse models where *Xbp1* is deleted specifically in Schwann cells, preventing XBP1s activation in these cells. We observed that *Xbp1* is dispensable for normal developmental myelination, myelin maintenance and remyelination after injury. However, *Xbp1* deletion dramatically worsens the hypomyelination and the electrophysiological and locomotor parameters observed in young and adult CMT1B neuropathic animals. RNAseq analysis suggested that XBP1s exerts its adaptive function in CMT1B mouse models in large part via the induction of ER proteostasis genes. Accordingly, the exacerbation of the neuropathy in *Xbp1* deficient mice was accompanied by upregulation of ER-stress pathways and of IRE1-mediated RIDD signaling in Schwann cells, suggesting that the activation of XBP1s via IRE1 plays a critical role in limiting mutant protein toxicity and that this toxicity cannot be compensated by other stress responses. Schwann cell specific overexpression of XBP1s partially re-established Schwann cell proteostasis and attenuated CMT1B severity in both the S63del and R98C mouse models. In addition, the selective, pharmacologic activation of IRE1α/XBP1 signaling ameliorated myelination in S63del dorsal root ganglia explants. Collectively, these data show that XBP1 has an essential adaptive role in different models of proteotoxic CMT1B neuropathy and suggest that activation of the IRE1α/XBP1 pathway may represent a therapeutic avenue in CMT1B and possibly for other neuropathies characterized by UPR activation.

## Introduction

In the peripheral nervous system (PNS) Schwann cells synthesize the myelin sheath that surrounds axons, allowing for fast conduction of the electrical impulses and protecting the axons from insults.^1^ Myelination requires Schwann cells to synthesize massive amounts of lipids and proteins that are then delivered to the plasma membrane through the secretory pathway.^2^ After the completion of myelination, the maintenance of myelin homeostasis is essential to preserve axon functionality.^3^ Myelin protein zero (MPZ/P0) is the most abundant protein of peripheral myelin, where it allows adhesion between adjacent myelin lamellae. Exact P0 folding and structure are essential for its function and for correct myelination.^4^ Indeed, single point mutations in *MPZ* are associated with autosomal dominant neuropathies with diverse phenotypes, ranging from classic demyelinating Charcot-Marie-Tooth disease type 1B (CMT1B), to the severe Dejerine-Sottas syndrome (DSS) and congenital hypomyelination,^5,6^ to primary axonal neuropathies also classified as CMT2J/I.^7^ The analysis of transgenic mice with authentic *Mpz* mutations has shown that they produce variable phenotypes through diverse gain of function mechanisms.^8^ Several mutant P0 proteins, among them S63del (CMT1B phenotype) and R98C (DSS phenotype), are retained in the endoplasmic reticulum (ER) where they induce ER stress and elicit an unfolded protein response (UPR).^9–11^

In mammalian cells, the UPR comprises three pathways activated downstream of the ER membrane proteins inositol requiring kinase 1α (IRE1α), activating transcription factor 6 (ATF6), and pancreatic ER kinase (PERK). These three proteins are activated in response to ER stress to initiate transcriptional and translational programs aimed at reducing the load of unfolded proteins through upregulation of chaperones, attenuation of protein synthesis and increased protein degradation.^12,13^ The importance of the UPR in myelin diseases, including CMT, has been demonstrated in several studies.^14,15^ We have provided genetic and pharmacological evidence that modulation of the PERK/P-eIF2α pathway can alter disease severity in rodent models of both CMT1B and CMT1A, the most common form of CMT, due to *PMP22* duplication. Ablation of the transcription factor CHOP (downstream of P-eIF2α) or inactivation of its target GADD34 from *MpzS63del* mice improved the CMT1B phenotype.^10,16^ Furthermore, Sephin1/IFB-088 a small molecule that prolongs translational attenuation by inhibiting P-eIF2α dephosphorylation, ameliorated disease parameters in models of CMT1B, CMT1A and amyotrophic lateral sclerosis in mice,^17,18^ strongly suggesting that the modulation of the UPR may represent a therapeutic option for CMT with altered proteostasis.

Among the UPR sensors, IRE1α is the most conserved through evolution.^12^ When misfolded proteins accumulate in the ER, IRE1α is activated through a mechanism involving oligomerization and autophosphorylation. This activates the IRE1α cytosolic endoribonuclease (RNase) activity. The IRE1α RNAse induces the unconventional splicing of X-box binding protein-1 (XBP1) mRNA leading to the production of spliced Xbp1 (XBP1s), a transcription factor that upregulates genes involved in ER biogenesis, protein folding and ER-associated degradation (ERAD).^19,20^ In conditions of ER-stress, IRE1α also leads to the degradation of a specific subset of mRNAs, in a process known as mRNA regulated IRE1-dependent decay (RIDD), which is thought to preserve ER homeostasis by reducing the load of cargo proteins entering the ER.^21^ XBP1 has a crucial role in mammalian physiology and diseases. Whole body *Xbp1* knock-out mice present hypoplastic foetal livers and die during early development due to anemia.^22^ Further, tissue-specific *Xbp1* conditional deletion causes severe abnormalities in plasma cells and pancreatic β-cells.^23,24^ Activation of XBP1s signalling was shown to be protective in several disease models, including Alzheimer’s disease^25^ and injury-induced retinal ganglion cell death.^26^ In contrast, XBP1s signalling was shown to be detrimental in models of amyotrophic lateral sclerosis,^27^ whereas contrasting results were obtained in Huntington disease models.^28,29^ The role of XBP1 in PNS development and neuropathy has however never been explored. To investigate this, we have generated and fully characterized from a clinical, neurophysiological, morphological and molecular perspective two different models of proteotoxic CMT1B in which *Xbp1* or *Xbp1s* where deleted or overexpressed, respectively, in a Schwann cells. Further, taking advantage of myelinating dorsal root ganglia (DRG) organotypic cultures, we have also explored how the pharmacologic modulation of of IRE1α/XBP1 signalling affect myelination in CMT1B.

## Materials and methods

### Animal models

All experiments involving animals were performed in accord with experimental protocols approved by the San Raffaele Scientific Institute Animal Care and Use Committee. *P0Cre*, *Mpz*S63del (*S63del* herein), *Mpz*R98C/+ (*R98C* herein), *Xbp1^f/f^* and *ROSA26*-*XBP1s* mice with relative genotyping procedures have been previously described.^8,9,30–32^ We initially crossed *S63del* and *P0Cre* mice with *Xbp1^f/f^* mice to obtain *S63del/Xbp1^f/+^* and *P0Cre/Xbp1^f/+^* mice that were then intercrossed to obtain the four experimental groups WT, *P0Cre/Xbp1^f/f^* (for the Schwann cell specific ablation of *Xbp1*, *Xbp1*^SC-KO^ mice herein), *S63del* and *S63del/P0Cre/Xbp1^f/f^* (*S63del/Xbp1^SC-KO^*herein). The same scheme was used to generate *R98C/Xbp1^SC-KO^* mice and the relative controls. Similarly, we crossed *P0Cre* and *ROSA26*-*XBP1s* mice to obtain *XBP1^SC-OE^* (Schwann cell specific overexpression of XBP1s) mice that were then crossed with either *S63del* or *R98C* mice to obtain *S63del/XBP1^SC-OE^*and *R98C/XBP1^SC-OE^* mice and the relative controls. *S63del* and *R98C* mice were maintained on the FVB/N background, whereas *P0Cre*, *Xbp1^f/f^* and *ROSA26*-*XBP1s* were on C57BL/6 genetic background. The analysis of all experimental genotypes was performed in F1 hybrid background FVB // C57BL6.

### Sciatic nerve crush

Adult WT and *Xbp1^SC-KO^* mice were anesthetized with tribromoethanol, 0.4 mg/gr of body weight and crush injury performed as described.^33^ Briefly, after skin incision, the sciatic nerve was exposed and crushed distal to the sciatic notch for 20 seconds with fine forceps previously cooled in dry ice. To identify the site of injury, forceps were previously dropped into vital carbon. The nerve was replaced under the muscle and the incision sutured. All the experiments have been performed on the distal portion of the crushed nerve.

### Behavioural analysis in Rotarod

Motor ability was assessed using the accelerating Rotarod (Ugo Basile, Comerio, Italy) as previously described.^16^ Briefly, adult transgenic and control littermates were tested in two sessions of three trials each per day (6h rest between the two daily sessions) for 3 consecutive days. During the test, the rod accelerated from 4 to 40 rotations per minute, and the time that the animal remained on the rod (maximum 900s) was measured.

### Neurophysiological analysis

The electrophysiological evaluation was performed with a specific EMG system (NeuroMep Micro, Neurosoft, Russia), as previously described.^10^ Mice were anesthetized with tribromoethanol, 0.02ml/g of body weight, and placed under a heating lamp to maintain constant body temperature. Sciatic nerve conduction velocity (NCV) was obtained by stimulating the nerve with steel monopolar needle electrodes. A pair of stimulating electrodes was inserted subcutaneously near the nerve at the ankle. A second pair of electrodes was placed at the sciatic notch to obtain two distinct sites of stimulation, proximal and distal along the nerve. Compound motor action potential (CMAP) was recorded with a pair of needle electrodes; the active electrode was inserted in muscles in the middle of the paw, whereas the reference was placed in the skin between the first and second digit. Sciatic nerve F-wave latency measurement was obtained by stimulating the nerve at the ankle and recording the responses in the paw muscle, with the same electrodes employed for the NCV study.

### Morphological and morphometric analysis

Transgenic mice were sacrificed at the indicated time points and sciatic nerves were dissected. Semi-thin section and electron microscope analyses of sciatic nerves, were performed as described^34^. The number of amyelinated or demyelinated axons were counted blind to genotype from post-natal day 30 (P30) or 6-month-old sciatic nerve semi-thin sections (0.5-1 μm thick) stained with tolouidine blue, in images taken with a 100x objective. We analyzed 1000–1800 fibers from a total of 10 - 20 fields per nerve (5 nerves). Ultrathin sections (90 nm thick) from P30 and 4 to 6-months transgenic and control sciatic nerves were cut using an ultracut ultramicrotome, stained with uranyl acetate and lead citrate and examined by electron microscopy (EM) (Leo 912 omega). G-ratio analysis (axonal diameter/fiber diameter) and the size distribution of myelinated fibers (based on axons diameter) were measured on digitalized non-overlapping electron micrograph images. 20–25 microscopic fields, randomly chosen, from nerves of 3–5 animals per genotype were analyzed.

### Dorsal root ganglia explant cultures

Dorsal-root-ganglia (DRG) were dissected from embryos at embryonic day 13.5 (E13.5) and plated singularly on collagen-coated coverslips as previously described.^16^ Myelination was induced with 50 μg/ml ascorbic acid (Sigma Aldrich). Treatment with IRE1α/XBP1 activating compounds (IXA) or with the IREα RNAse inhibitor 4μ8c at the indicated concentration was applied for 2-weeks in parallel to the induction of myelination. Samples were then fixed and rat anti-MBP (1/5), and rabbit anti-NF-H (1/1000, EMD Millipore) primary antibodies were added o/n at 4°C. The following day, DRGs were washed and FITC- or TRITC-conjugated secondary antibodies (1:200, Cappel) were added for 1h at room temperature. Specimen were incubated with DAPI (1:1000, SIGMA) and mounted with VectaShield (Vector Laboratories). 8-10 images were taken from each DRG using a fluorescence microscope (Leica DM5000) with a 20x objective, and the number of MBP+ internodes in each image was counted.

### Protein extraction and Western blot

Sciatic nerves were dissected from mice at the indicated age and frozen in liquid nitrogen. Frozen nerves were pulverized on dry ice and proteins were extracted in denaturing lysis buffer (Tris-HCl 50 mM pH 7,5, NaCl 150 mM, EDTA 10 mM, 2% SDS) containing protease inhibitor cocktail (PIC 100X, Roche) and phosphatase inhibitors (Roche). Total protein concentration was determined by BCA assay (Pierce) following manufacturer’s instructions. Equal amounts of proteins were separated by SDS-PAGE (Biorad) and gels were transferred onto nitrocellulose membrane (GE Healthcare). Membranes were blocked with 5% milk (milk powder/ 1X PBS-Tween 0,1%) or 5% BSA (BSA powder/ 1X PBS-Tween 0,1%) and incubated with primary antibodies diluted in 1% milk or 1% BSA/ 1X PBS-Tween 0,1% at 4°C overnight. HRP-conjugated antibodies were diluted in 1% milk or 1% BSA/ 1X PBS-Tween 0,1% and incubated 1h at room temperature. Signals were detected by ECL method. Densitometric analysis was performed with NIH-Image-J software. The following primary antibodies were used: mouse anti β-tubulin (T4026, Sigma), mouse anti-MBP (MAB 382, Chemicon), rabbit anti-PMP22 (AB2011052 Abcam), rabbit anti-P-eIF2α (Ser51)(D9G8) XP and anti-eIf2α (D7D3) XP (Cell Signaling), rabbit anti-BiP (NB300520, Novus Biologicals), rat anti-GRP94 (ab2791, Abcam), rabbit anti-IRE1a (3294, Cell Signaling), rabbit anti P-JNK and JNK (4668 and 9252, Cell Signaling), chicken anti-P0 (PZO, Aves). Peroxidase-conjugated secondary antibodies (anti-rabbit HRP, DAKO, P0448; anti-chicken IgG-peroxidase, Sigma; anti-mouse IgG peroxidase, A3682, Sigma) were visualized using Amersham ECL or ECL Prime 225 reagent (GE Healthcare) for high-sensitivity chemiluminescent protein detection with Uvitec gel analysis systems or using enhanced chemiluminescence (ECL) reagents (Bio-Rad) with autoradiography film (Kodak Scientific Imaging Film, Blue XB). Total proteins were visualized via staining with Coomassie Brilliant blue R250 staining solution (Bio-Rad). Densitometric quantification was performed with ImageJ.

### mRNA extraction and real-time qPCR

Total RNA was extracted with Trizol (Roche Diagnostic GmbH, Germany) and retrotranscribed as previously described.^16^ cDNAs were quantified by quantitative real-time PCR on a MX3000 apparatus (Stratagene, La Jolla, CA, USA) by using specific primers. Primer sequences are provided in Supplementary table 1. PCR amplification was performed in a volume of 20 μl containing 5 ng cDNA, 300 nM of each primer, GoTaq qPCR Master Mix (Promega, Madison, WI, USA). Gene expression changes were normalised to 36B4 or Ppia gene expression by using the ΔΔCT method.

### RNA-sequencing

At least 1000 ng total RNA was purified from P30 sciatic nerves from WT, *Xbp1^SC-KO^, S63del and S63del/Xbp1^SC-KO^* mice using the Zymo RNA Clean & Concentrator-5 (#R1013) and sent to Genewiz (South Plainfield, NJ) for library preparation after PolyA selection and Illumina sequencing (Illumina HiSeq 2x150bp). Illumina sequencing data were mapped to the GRCm38/mm10 genome using the STAR aligner.^35^ Data were analyzed using DESeq2 to determine differentially regulated genes (p-value < 0,5).^36^ RNA-seq data are deposited in NCBI GEO under accession number GSE252089.

### Statistical analysis

Sample size was not predetermined with any statistical method, but our sample size is similar to that generally used in the field. Graphs and data were analyzed using GraphPad Prism Software. Data show the mean ± Standard Error of Mean (SEM). Unpaired, 2 tails, Student’s *t* test (for comparison between two groups) or ANOVA (for comparison between three or four groups) with Tukey post-hoc multiple comparison test were used as specified in the figure legends; significance levels (*P* values) were marked on figures as follows: \**P* ≤ 0.05, \*\**P* ≤ 0.01, \*\*\**P* ≤ 0.001, \*\*\*\**P* ≤ 0.0001.

## Results

### *Xbp1* is dispensable for peripheral myelin formation, maintenance and remyelination after injury

To evaluate the role of XBP1 in nerve development and CMT1B pathogenesis, we generated mice with specific ablation of *Xbp1* in Schwann cells (*Xbp1^SC-KO^* mice) by crossing *Xbp1* floxed mice^31^ with *P0Cre* mice.^30^ *Xbp1^SC-KO^* mice were then crossed with *S63del*-CMT1B mice^8^ to obtain *S63del/Xbp1^SC-KO^* and the respective P0Cre-negative controls. To test the efficiency and specificity of P0Cre-mediated recombination, we performed PCR reactions on genomic DNA extracted from 1-month-old sciatic nerves and several other tissues (**Supplementary Fig. 1A**). In sciatic nerves, the recombined *Xbp1* KO (350bp) band was specifically detected in P0Cre-positive samples, but not in P0Cre negative control nerves, as expected. Moreover, in *Xbp1^SC-KO^* mice, the recombination band appeared only in sciatic nerve and not in other tissues. Accordingly, the mRNA levels of *Xbp1*, which were increased in *S63del* nerves due to the ongoing UPR, were largely reduced in nerve extracts from *Xbp1^SC-KO^*and *S63del/Xbp1^SC-KO^* mice (**Fig. 1A**).

**Figure 1.**
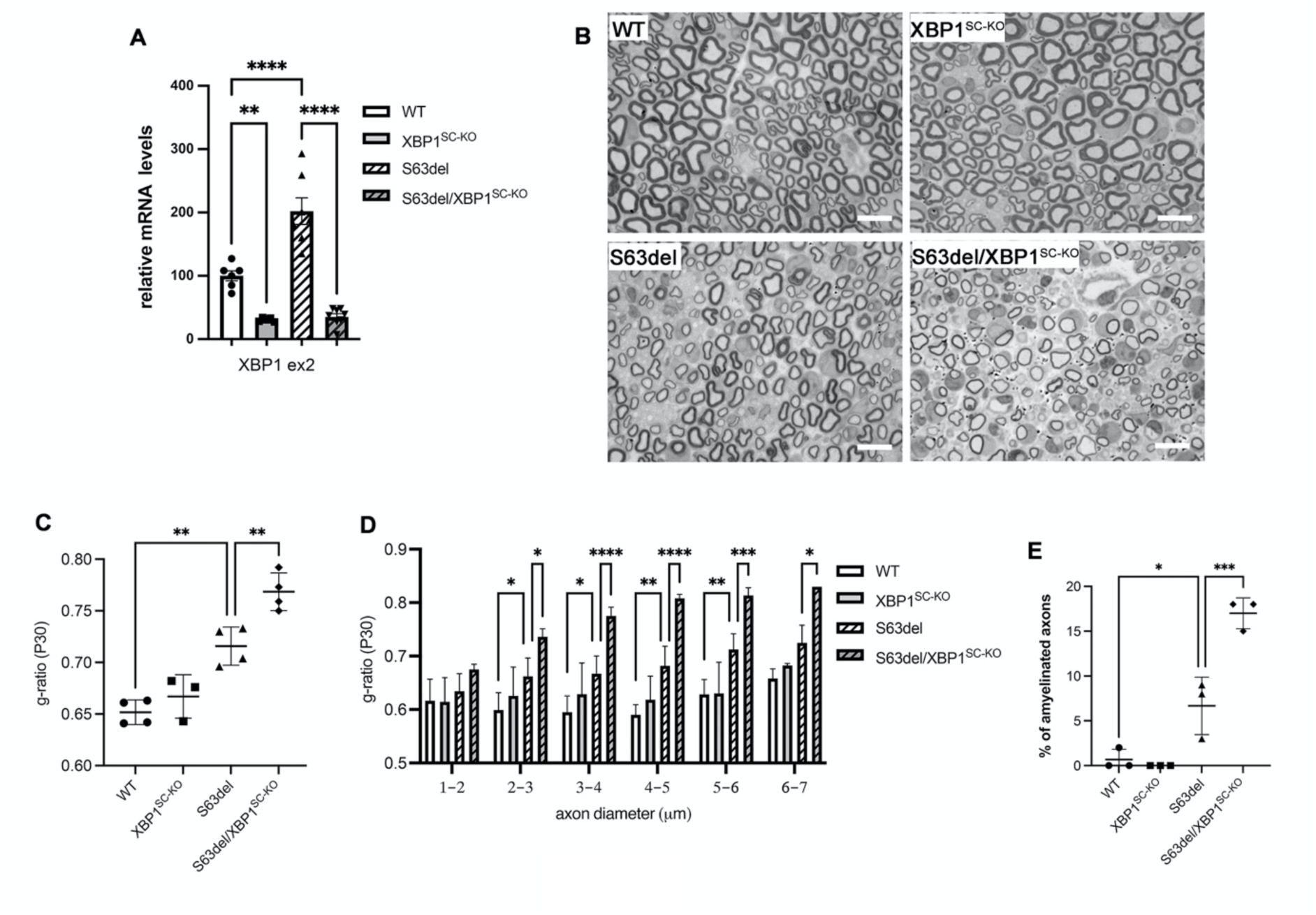
Schwann cell specific ablation of *Xbp1* worsens pathology in developing *S63del* mice. (**A**) qRT-PCR analysis for *Xbp1* exon 2 (which is deleted following CRE mediated recombination) on mRNA extracts from P30 sciatic nerves; *n* = 5-7 reverse transcription (RT) per genotype from independent nerves. (**B**) Transverse semithin sections from P30 sciatic nerves. Scale bar 10μm. (**C**) G-ratio analysis performed on P30 sciatic nerves semithin sections; *n* = 3-4 mice per genotype (**D**) Average g-ratio plotted by axon diameter. (**E**) Percentage of amyelinated axons (axons > 1μm in a 1:1 relationship with a Schwann cell but not myelinated); 8-10 microscopy field from semithin sections per mouse were evaluated from *n* = 3 mice per genotype. Error bars represent SEM and \*\*\*\**P* < 0.0001, *** *P* < 0.001, ** *P* <0.01, * *P* <0.05 by one-way ANOVA followed by Tukey post hoc test.

Given the essential role of XBP1 for the differentiation and function of several highly secretory cells such as plasma cells, β-cells and adipocytes,^23,24,37^ we reasoned that the ablation of *Xbp1* could have detrimental effects on Schwann cells differentiation and myelination. To test this hypothesis, we examined *Xbp1^SC-KO^* sciatic nerve morphology in postnatal day 5 (P5) (**Supplementary Fig. 1C)**, P30 (**Fig. 1B**), P180 (**Fig. 2D**) and in 1-year-old animals (**Supplementary Fig. 1D**). At all these time points, *Xbp1^SC-KO^* nerves did not display any gross myelin abnormality. At P30 and P180, myelin thickness was normal as measured by morphometric g-ratio analysis on semithin sections (**Fig. 1C and Fig. 2G**), and the axon size distribution was unchanged as compared to WT controls (data not shown), suggesting that XBP1/XBP1s is not required for myelination.

**Figure 2.**
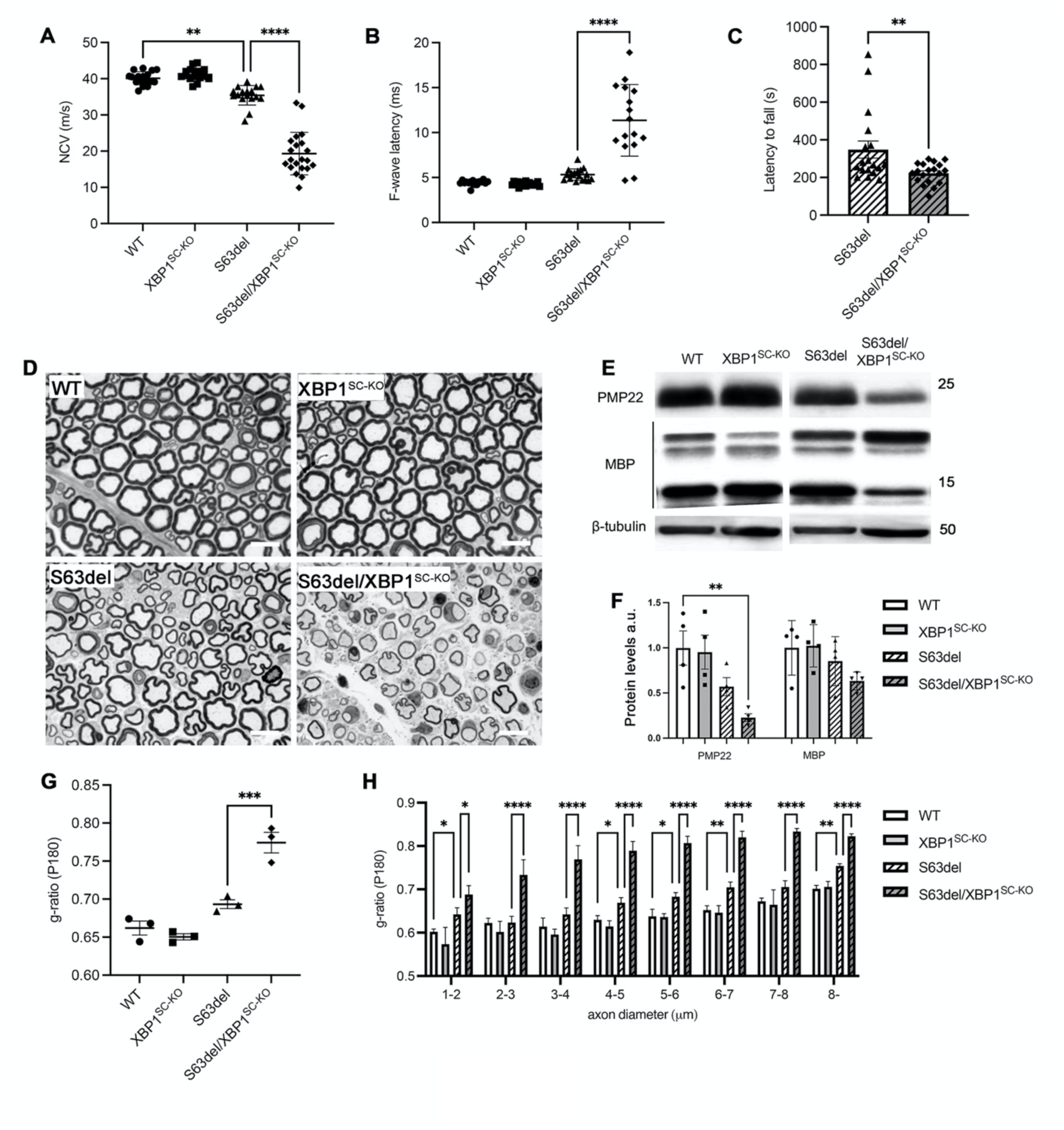
Schwann cell specific ablation of *Xbp1* worsens neurophysiological, behavioral and morphological parameters in adult *S63del* mice. Analysis of nerve conduction velocities (NCV) (**A**) and F-wave latencies (**B**) in 6-months old mice; *S63del/Xbp1^SC-KO^* mice show severe neurophysiological worsening as compared to *S63del*; n = 12-16 mice per genotype. \*\*\*\**P* < 0.0001 and ** *P* <0.01 by one-way ANOVA followed by Tukey post hoc test. (**C**) Rotarod analysis performed at 6 months show reduced motor capacity in *S63de/Xbp1^SC-KO^* mice. ** *P* <0.01 by unpaired Student’s *t*-test. (**D**) Transverse semithin sections from 6-months old sciatic nerves. Scale bar 10μm. (**E**) Western blot analysis on sciatic nerve extracts and (**F**) relative quantification for the myelin proteins MBP and PMP22; β-tubulin was used as loading control. ** *P* <0.01 by one-way ANOVA followed by Tukey post hoc test; *n* = 4. (**G**) G-ratio analysis performed on 6-month-old sciatic nerve semithin sections and (**H**) average g-ratio plotted by axon diameter from *n* = 3 mice per genotype. Error bars represent SEM and \*\*\*\**P* < 0.0001, *** *P* < 0.001, ** *P* <0.01, * *P* <0.05 by one-way ANOVA followed by Tukey post hoc test.

Recent work showed activation of ER-stress in the distal segment of injured sciatic nerve and suggested that *Xbp1* deficiency in the nervous system decreased axonal regeneration after injury.^38^ To assess whether Schwann cell specific XBP1 was required for remyelination after nerve injury, we performed sciatic nerve crush experiments on 2-month-old *Xbp1^SC-KO^*mice and WT controls and looked at the extent of remyelination 20 days after injury (**Supplementary Fig. 2**). Morphological and morphometric analysis performed on semithin sections did not reveal any significant difference in the extent of remyelination, with comparable numbers of remyelinated fibres in *Xbp1^SC-KO^* and WT injured controls. Taken altogether, these data show that *Xbp1* is dispensable for Schwann cell developmental myelination, for myelin maintenance in the adult and for remyelination after injury.

### Ablation of *Xbp1* worsens pathological defects in S63del CMT1B mice

We next investigated the role of XBP1 in the *S63del* CMT1B model. In *S63del* nerves, the IRE1α/XBP1 pathway is activated as early as P5 and remains activated throughout adulthood, as shown by qRT-PCRs for spliced *Xbp1* (*Xbp1s*) (**Supplementary Fig. 1B**). At the morphological level, genetic ablation of *Xbp1* in Schwann appeared to already aggravate the neuropathic phenotype at P5 (**Supplementary Fig. 1C**). By P30, a time point that roughly corresponds to the peak of myelination and ER-stress in *S63del* mice,^16^ *S63del/Xbp1^SC-KO^*sciatic nerves appeared severely compromised, as shown by images of semithin sections (**Fig. 1B**). Accordingly, we observed an increase in the mean g-ratio in *S63del/Xbp1^SC-KO^* nerves as compared to *S63del*, that was consistent for all axon sizes (**Fig. 1D**). In addition, we observed a significant increase in frequency of amyelinated axons (axons with diameter >1mm blocked in a 1:1 relationship with a Schwann cell) in *S63del/Xbp1^SC-KO^*nerves (**Fig. 1C**) Electron microscopy (EM) analysis performed on P30 sciatic nerves confirmed equal myelination in *Xbp1^SC-KO^* and WT nerves. Conversely, the EM analysis showed hypomyelination and increase in amyelinated axons in *S63del/Xbp1^SC-KO^* nerves as compared to the *S63del* control, without significant alterations in Schwann cell morphology (**Supplementary Fig. 3A**).

Adult *S63del* mice manifest altered neurophysiology, locomotor impairment and progressive demyelination, similar to the alterations found in patients.^8,39^ To test how the genetic ablation of *Xbp1* would affect these features, we performed neurophysiological analysis on mice at 6 months of age. We confirmed the alteration of NCV in *S63del* mice compared with control mice;^8^ *S63del/Xbp1^SC-KO^* showed a very severe impairment of NCV as well as F-wave latencies (**Fig. 2A-B**). *S63del/Xbp1^SC-KO^* mice also showed a reduced latency to fall on the rotarod test as compared to *S63del* (**Fig. 2C**).

In line with neurophysiological and behavioural findings, morphological analysis of 6-month and 1-year old nerves confirmed the hypomyelination of *S63del* nerves as compared to WT and an extensive worsening of the phenotype in *S63del/Xbp1^SC-KO^* (**Fig. 2D and Supplementary Fig. 1D**) as corroborated by g-ratio measurements (**Figure 2G-H**). Finally, western blot (WB) analysis for two of the major myelin components, myelin basic protein (MBP) and peripheral myelin protein 22 (PMP22) confirmed the strong reduction of bulk myelin in 6-month-old *S63del/Xbp1^SC-KO^* nerves (**Fig. 2E-F**).

### Ablation of *Xbp1* worsens the pathology of R98C mice

To better understand the relevance of our findings for CMT1B pathogenesis, we then asked whether the protective role of XBP1 could be replicated in a second model of CMT1B that shows high levels of UPR activity, the *Mpz*-*R98C* mouse.^9^ First, we evaluated the morphology of *R98C* and *R98C/Xbp1^SC-KO^* nerves at P30 and found that overall g-ratio was only marginally increased in *R98C/Xbp1^SC-KO^*as compared to *R98C* (0.78 ± 0.01 vs 0.80 ± 0.01, *P* = 0.3), (**Supplementary Fig 3B-C)**. However, by 6-months of age neurophysiological analysis showed that NCV and F-wave latency, already largely compromised in *R98C* nerves, were further worsened in *R98C/Xbp1^SC-KO^* mice (**Fig. 3A-B**), that also displayed reduced motor abilities as compared to *R98C* mice, as measured on the accelerating rotarod (**Fig 3C**). Accordingly, morphological and morphometric analysis performed on both semithin and ultrathin EM sections (**Fig. 3D and 3E**) showed a worsening of the dysmyelinating phenotype of *R98C* mice following Schwann cell specific ablation of *Xbp1*, as confirmed by a significant increase in g-ratio (**Fig. 3F**), with small and medium calibre axons (up to 7μm) more affected (**Fig. 3G**). Altogether these results indicate that the activation of the IRE1α/XBP1 pathway of the UPR represents a protective event in early and late stages of different models of proteotoxic CMT1B.

**Figure 3.**
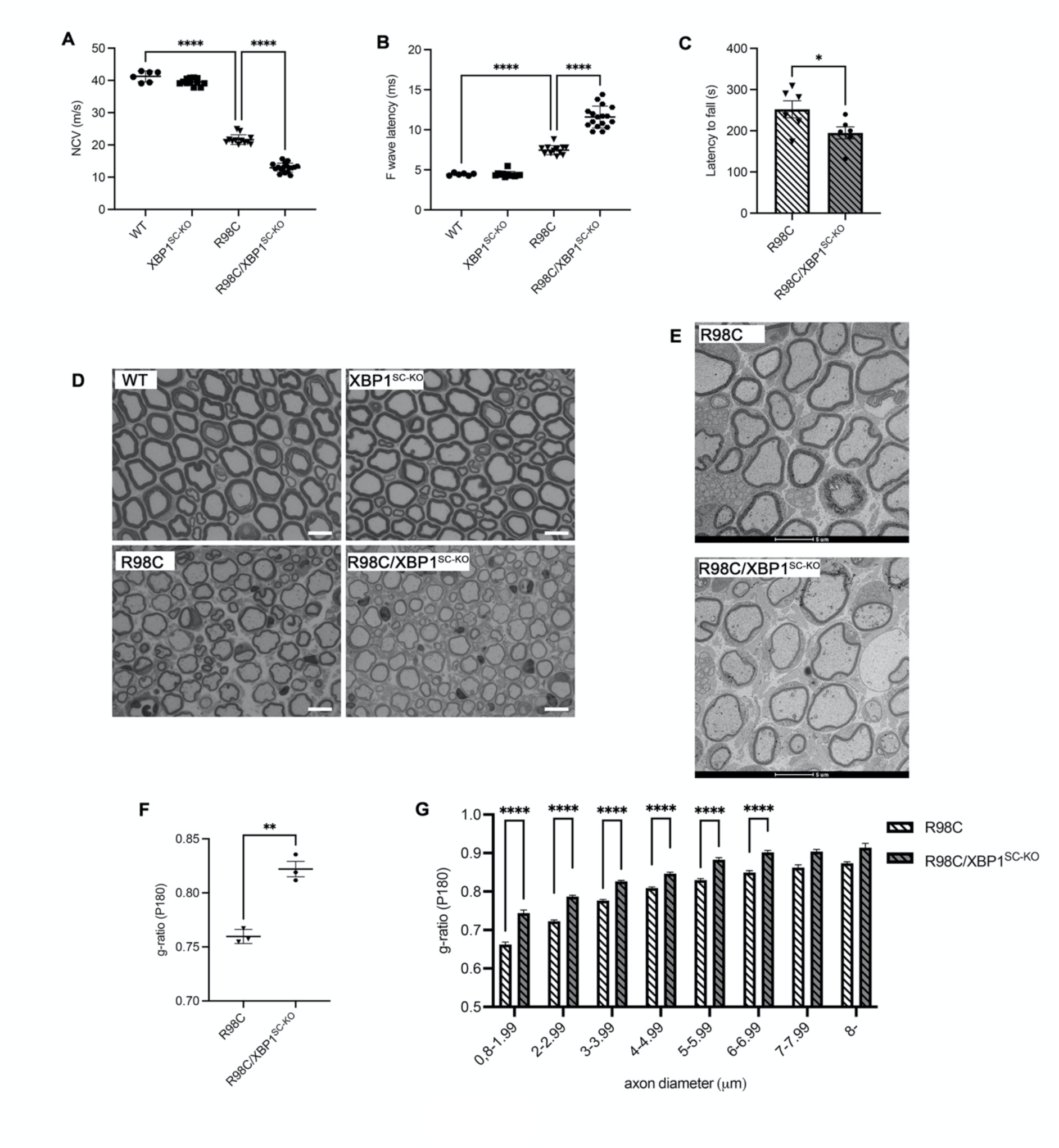
Schwann cell specific ablation of XBP1 worsens neurophysiological, behavioral and morphological parameters in adult *R98C* mice. Analysis of nerve conduction velocities (NCV) (**A**) and F-wave latencies (**B**) in 6-months old mice. n = 6-12 mice per genotype. \*\*\*\**P* < 0.0001 by one-way ANOVA followed by Tukey post hoc test. (**C**) Rotarod analysis performed at 6-months show reduced motor capacity in *R98C/Xbp1^SC-KO^* mice. * *P* <0.05 by unpaired Student’s *t*-test. (**D**) Transverse semithin sections from 6-months old sciatic nerves. Scale bar 10μm. (**E**) Electron microscopy on sciatic nerve transverse sections from *R98C* and *R98C/Xbp1^SC-KO^* mice at 6-months show severely reduced myelin thickness in CMT1B mice lacking *Xbp1*; Scale bar 5μm. (**F**) G-ratio analysis performed on 6-month-old sciatic nerve semithin sections and (**G**) average g-ratio plotted by axon diameter from *n* = 3 mice per genotype. Error bars represent SEM and \*\*\*\**P* < 0.0001, ** *P* <0.01, * *P* <0.05 by unpaired Student’s *t*-test.

### *Xbp1* ablation limits UPR and ERAD activation, exacerbates the ER-stress response and impairs Schwann cell differentiation in S63del CMT1B nerves

To gain insight in the molecular mechanisms underlying the adaptive role of XBP1 in Schwann cells under proteotoxic ER-stress, we performed RNA sequencing on mRNA extracted from P30 *S63del* and *S63del/Xbp1^SC-KO^* sciatic nerves, with WT and *Xbp1^SC-KO^* nerves serving as controls. As already observed from nerve morphology and neurophysiology, XBP1 deletion in healthy mice had no major influence on nerve transcriptome with fewer than 150 genes significantly up- or down-regulated in *Xbp1^SC-KO^* nerves as compared to WT. Conversely, in *S63del* nerves 4585 genes (of which 1637 with a fold change > 1.5) were found to be significantly, differentially expressed compared to WT. This number increased to a total of 8155 (4640 with fold change > 1.5) in the double mutant *S63del/Xbp1^SC-KO^*, indicating that the ablation of *Xbp1* dramatically impacts the pool of genes found to be dysregulated in CMT1B nerves. Accordingly, volcano plots highlighted how the significance and fold change with which up and downregulated genes in *S63del/Xbp1^SC-KO^* differ from WT is largely increased as compared to S63del, confirming a vast alteration in the transcriptional landscape (**Fig. 4A-B**).

**Figure 4.**
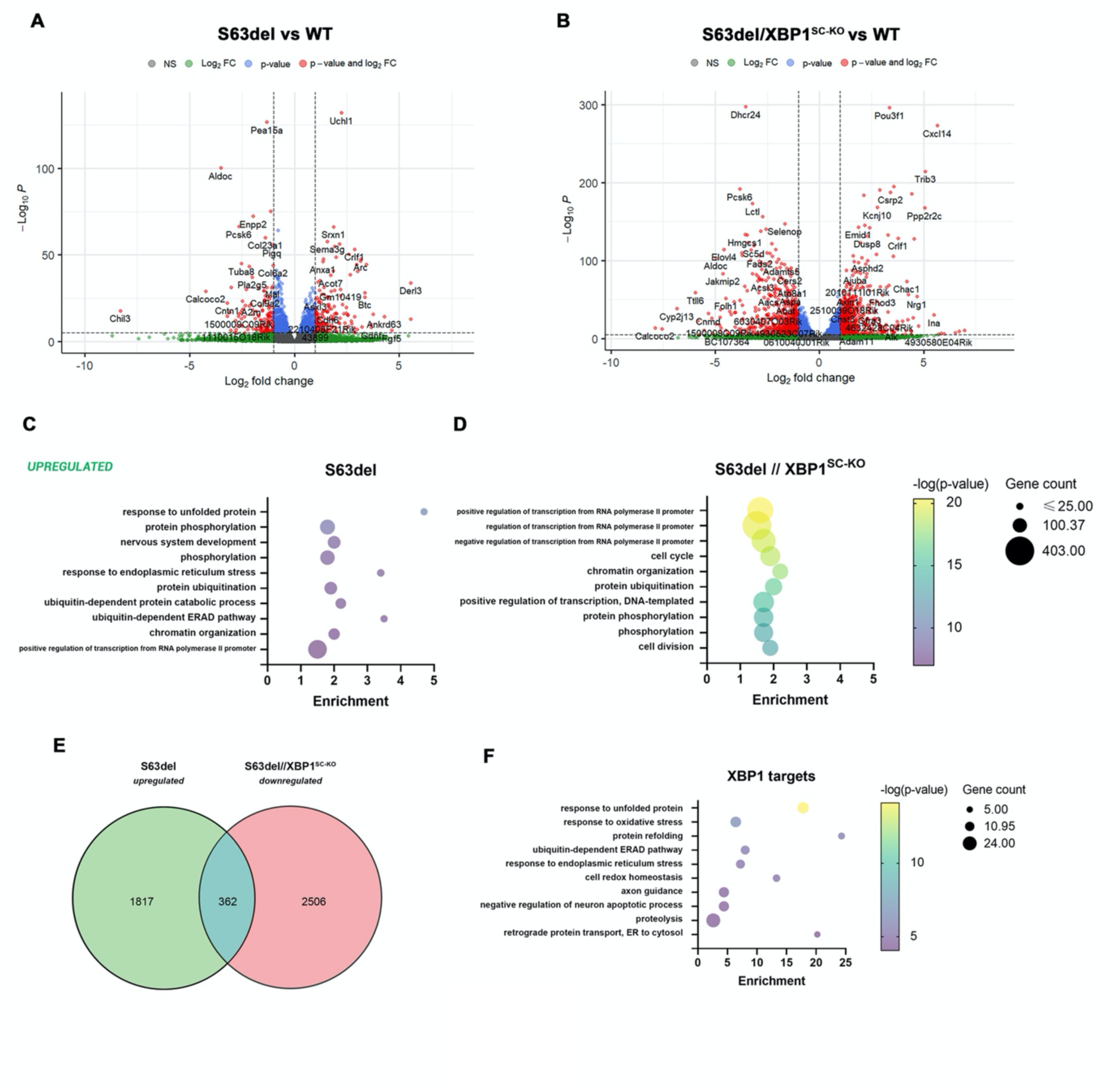
RNA-sequencing in *S63del* and *S63de/Xbp1^SC-KO^*. (**A**) Volcano plot showing the genes differentially regulated in P30 *S63del* nerves in comparison to WT. (**B**) Volcano plot showing the genes differentially regulated in P30 *S63del/Xbp1^SC-KO^* nerves in comparison to WT. (**C**) Gene ontology (GO) of biological processes upregulated in *S63del* vs WT. (**D**) GO of biological processes upregulated in *S63del/Xbp1^SC-KO^* vs WT. (**E**) Venn-diagram of genes upregulated in *S63del* nerves as compared to WT, and genes down-regulated in *S63del/Xbp1^SC-KO^* as compared to *S63del*. The 362 common genes are likely direct or indirect targets of XBP1. (**F**) GO of the 362 putative targets of XBP1s.

Gene Ontology (GO) annotation performed on the upregulated genes, identified “response to unfolded protein”, “response to ER stress” and “protein degradation (via the ubiquitin-ERAD pathway)” among the most significant and enriched biological processes in *S63del* nerves (**Fig. 4C**), in line with what was observed in previous microarray studies.^16^ Conversely, in *S63del/Xbp1^SC-KO^* nerves GO processes “UPR” and “ERAD” were no longer in the top hits, where instead we found processes related to “regulation of transcription” and “cell cycle” (**Fig. 4D**). Of note, the GO terms related to the ER-stress response were still present amongst the upregulated processes in *S63del/Xbp1^SC-KO^* nerves but were generally less enriched (**Supplementary Fig. 4A**). The genes that belonged to these categories in *S63del/Xbp1^SC-KO^*were mostly known targets of the PERK/ATF4 arm of the UPR (e.g., *Chop/Ddit3*, *Asns*, *Trib3* and *Mthfd2*), and the ATF6 pathway, (*BiP/Hspa5*, *Grp94* and *Sel1l*). Intriguingly, most of the PERK/ATF4 targets were increased in *S63del/XBP1^SC-KO^* nerves as compared to *S63del* (for example, compared to WT nerves, *Chop/Ddit3*, *Asns*, and *Mthfd2* where increased 1.9, 2.1, 1.6 fold and 6.9, 7.2 and 12-fold in *S63del* and *S63del/Xbp1^SC-KO^* respectively) whereas the ATF6 targets remained basically unchanged. This suggests that in the absence of XBP1s, stress pathways (in particular downstream of the PERK branch) are hyperactivated but do not appear to be able to compensate for the lack of XBP1s. As mentioned, regulation of transcription and cell cycle were instead the top GO hits in *S63del/Xbp1^SC-KO^* nerves. In this category we identified a series of early transcription factors, such as *Id2, Sox2, Pou3f1 and c-Jun*, that can act as negative regulators of myelination.^40–42^ These factors have been shown to be increased in many neuropathic models, including *S63del*,^41,43^ and appear to be further increased in the absence of *Xbp1*, supporting the morphological finding of exacerbated dysmyelination in *S63del/Xbp1^SC-KO^* nerves.

On the other hand, downregulated genes in *S63del* nerves were mostly associated with “lipid/cholesterol metabolic processes” or “cell adhesion”and these terms remained the most relevant when XBP1s was removed (**Supplementary Fig. 4B**).

The RNAseq analysis thus suggested that XBP1s is mostly involved in transcriptional activation of target genes. Hence, we reasoned that potential XBP1s targets (direct or indirect) should be activated in *S63del* but not in *S63del/Xbp1^SC-KO^* as compared to WT, and as a result they may appear downregulated in *S63del/Xbp1^SC-KO^ vs S63del*. This approach identified a total of 362 genes whose activation in *S63del* was lost upon *Xbp1* removal (**Figure 4E**). This set of genes was subjected to GO analysis. Unsurprisingly, this showed that most XBP1s targets are related to the “response to unfolded protein and oxidative stress”, and to “protein refolding or degradation via ERAD” (**Figure 4F**). We therefore refined our analysis performing a GO analysis for *“cellular compartment”* and focused our attention on genes that were annotated with *“endoplasmic reticulum”*. Among them were genes who had previously been described as components of ERAD and proteasomal degradation (*Derl3, Erdj4/Dnajb9, Erdj6/Dnajc3* and *Uchl1*), and targets of the UPR and/or ER-stress response pathways (*Sdf2l1, Hyou1 and Creld2*). Taken together, these results show that XBP1 is protective in the S63del neuropathy and suggest that its activation likely reduces ER stress through the stimulation of pro-survival events of the UPR, including ER proteostasis.

To validate these findings, we first performed real time qPCR analysis on mRNA extracted from an independent set of P30 sciatic nerves. This analysis confirmed that *Erdj4* and *Erdj6*, two ER co-chaperones known to associate to and promote the degradation of misfolded proteins, were upregulated in *S63del* nerves, but no longer increased in the absence of XBP1s (**Fig. 5A**). Next, we measured the levels of activation of the other UPR pathways, by evaluating the mRNA levels of PERK pathway targets *Chop*, *Gadd34*/*Ppp1r15a* and *Mthfd2* and of ATF6 targets *BiP*, *Grp94* and *Sel1l*; this analysis confirmed an hyperactivation of the PERK pathway in S63del nerves deleted of *Xbp1*, whereas the ATF6 pathway appeared mostly unchanged (**Fig 5B-C**). This is consistent with what was shown by RNAseq. Finally, we looked at the expression levels of the negative regulator of myelination *Id2*, which was confirmed to be increased in *S63del* nerves as compared to WT and was further increased after *Xbp1* ablation (**Fig. 5D**). Overall, these data confirmed that as early as P30, *Xbp1* ablation in S63del nerves exacerbates ER-stress and hampers Schwann cells differentiation.

**Figure 5.**
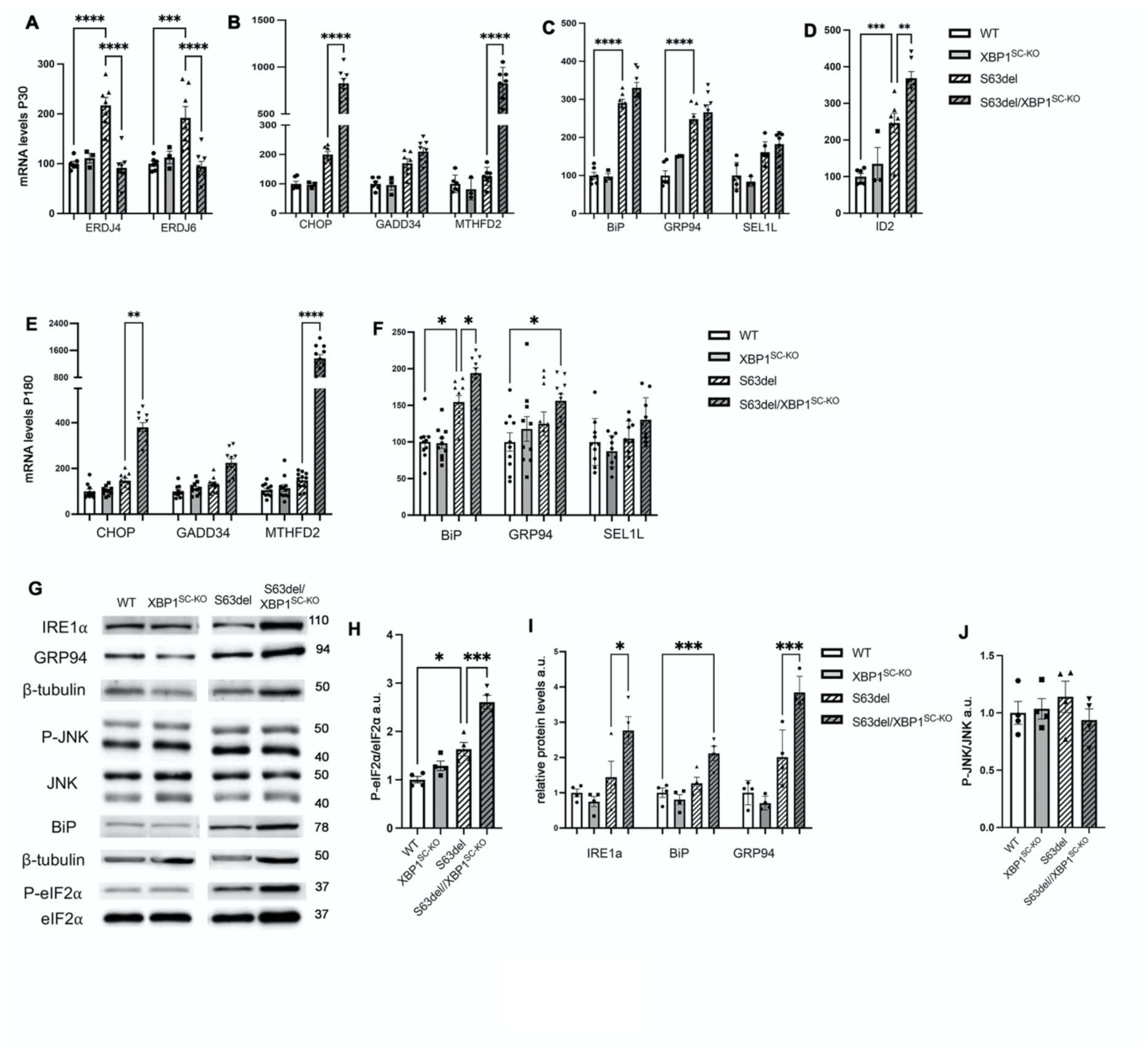
Ablation of *Xbp1* limits ER-associated degradation, exacerbates ER-stress and impairs Schwann cell differentiation in *S63del* mice. qRT-PCR analysis on mRNAs extracted from P30 sciatic nerves for (**A**) the XBP1s targets *Erdj4* and *Erdj6* (**B**) the PERK/ATF4 pathway targets *Chop, Gadd34* and *Mthfd2* and (**C**) the ATF6 pathway targets *BiP, Grp94* and *Sel1l*. (**D**) qRT-PCR analysis for the negative regulator of myelination *Id2*. (**E-F**) qRT-PCR analysis on mRNAs extracted from P180 sciatic nerves for the (**E**) PERK/ATF4 targets *Chop, Gadd34* and *Mthfd2* and (**F**) the ATF6 pathway targets *BiP, Grp94* and *Sel1l*. For all qRT-PCR experiments *n* = 8-10 RTs from independent nerves for each genotype. Values are expressed as fold change relative to WT (arbitrarily set to 100); error bars represent SEM. (**G**) Representative Western blots form P180 sciatic nerve extracts for IRE1a, GRP94, P-JNK, BiP and P-eIF2a; b-tubulin was used as loading control. One representative blot of four (quantified in the graphs in **H, I** and **J**) is shown per each genotype. Values are expressed as arbitrary units relative to WT that was set to 1. Bars represent SEM. For all panels \*\*\*\**P* < 0.0001, *** *P* < 0.001, ** *P* <0.01, * *P* <0.05 by one-way ANOVA followed by Tukey post hoc test.

The hyperactivation of the PERK pathway in *S63del/Xbp1^SC-KO^* was confirmed at P180, as shown by increased expression of PERK target genes (**Fig 5E**). Similarly *BiP*, the main target of the ATF6 pathway and the key regulator of the ER-stress response, was significantly increased (**Fig. 5F**). To further corroborate these findings, we also checked the protein levels of a subset of UPR factors and observed increased protein levels for P-eIF2α (PERK pathway) **(Fig. 5G-H)**, as well as BiP and GRP94 (ATF6 pathway) in *S63del/Xbp1^SC-KO^* nerves relative to *S63del* (**Fig. 5G and I).** Importantly, we detected a comparable increase in ER-stress levels and early Schwann cell transcription factors also in sciatic nerves from *R98C/Xbp1^SC-KO^* mice (data not shown), suggesting a common pathomechanism following XBP1 ablation.

### IRE1 and RIDD activation do not contribute to worsening the CMT1B phenotype after *Xbp1* ablation

The WB analysis performed in P180 nerves also detected increased levels of IRE1α **(Fig 5G**). Previous works suggested that genetic ablation of *Xbp1* results in hyperactivation of IRE1α RNAse activity.^32,44,45^ Accordingly, we found that in *S63del/Xbp1^SC-KO^* the mutant *Xbp1* mRNA (lacking exon2 and so not encoding a functional protein) was spliced at a substantially higher level than in *S63del* nerves (**Supplementary Fig. 5A).** In addition to its role in the splicing of *Xbp1* mRNA, active IRE1α can promote cell death via activation of the JNK pathway downstream of phosphorylated IRE1α,^46^ and induce a pathway known as regulated IRE1α dependent decay (RIDD) that leads to preferential degradation of ER-localized mRNAs by the IRE1α RNAse. This RIDD activity is thought to function by reducing the amount of proteins entering the ER; however, in response to unmitigated ER stress, RIDD can also initiate apoptosis.^21,47,48^ As assessed by the phosphorylation of JNK, the JNK pathway was however not activated either by S63del stress or by *Xbp1* deficiency (**Fig 5G and J**). On the other hand, we observed a decrease in mRNA levels for the known RIDD target *Blos1* in P30 and P180 *S63del/Xbp1^SC-KO^* nerves compared to *S63del*, indicating that RIDD is hyperactivated in the nerves lacking *Xbp1* (**Supplementary Figure 5B-C**). This further confirms an overall increase in IRE1α RNAse activity.

This hyperactivation of RIDD raised the question of whether the worsening of the phenotype observed in in *S63del/Xbp1^SC-KO^* nerves was solely due to loss of XBP1s or if excessive IRE1α RIDD activity could also contribute. To answer this question, we turned to a simpler *ex-vivo* myelination system, and prepared dorsal root ganglia (DRG) cultures from WT, *Xbp1^SC-KO^, S63del* and *S63del/Xbp1^SC-KO^*E13.5 embryos. As observed in previous reports,^16,18^ myelination is largely reduced in *S63del* cultures compared to WT (**Supplementary Fig 5D**). Consistent with the *in vivo* data, the genetic ablation of *Xbp1* does not affect myelination in DRG expressing WT P0 but further reduces the capacity of Schwann cells to produce myelin in cultures expressing the P0S63del mutant (**Supplementary Fig 5D-F**). To test if this was caused only by the lack of XBP1s or by IRE1α RIDD hyperactivation, we treated WT and *S63del* DRGs with 4μ8c (**Supplementary Fig 5G)**, a specific non-competitive inhibitor of the endoribonuclease activity of IRE1α.^49^ The treatment was effective in almost completely abolishing IRE1α RNAse activity as measured by *Xbp1s* levels (**Supplementary Fig. 5H**). If hyper-induction of IRE1α (and thus RIDD) was playing a detrimental role, *S63del* cultures treated with 4μ8c should show improved myelination as compared to *S63del/Xbp1^SC-KO^* DRGs. Strikingly however, treatment of S63del DRG explants with 4μ8c reduced myelination at least as much as *Xbp1* deletion (compare **Supplementary Fig. 5F and 5I)** without impairing myelination in WT controls (**Supplementary Fig. 5G)**. These results suggest that the ablation of *Xbp1 per se* is the main factor in the severely worsened CMT1B defects.

### Genetic overexpression of XBP1s ameliorates CMT1B neuropathies *in vivo*

Overall, our data indicated an adaptive role for XBP1 in proteotoxic CMT1B. Interestingly, in both *S63del* and *R98C* nerves only a small proportion of the available XBP1 is spliced.^9,10^ We thus wondered if further increase in XBP1s levels in neuropathic Schwann cells would attenuate the disease. To test this hypothesis, we crossed *ROSA-XBP1s* mice, in which a loxP flanked transcriptional STOP sequence coupled to the human XBP1s (*hXBP1s*) cDNA was inserted into the ROSA26 locus,^32^ with *P0Cre* and then *S63del* mice, to obtain *XBP1^SC-OE^* and *S63del/XBP1^SC-OE^* mice, in which Cre activation resulted in hXBP1s expression only in Schwann cells. Consistently, we observed that the levels of the XBP1s target gene *Erdj4* were increased in *S63del/XBP1^SC-OE^*as compared to *S63del* at P30, confirming a further activation of the pathway (**Fig. 6A**). Morphological and morphometric analysis, performed on P30 sciatic nerve semithin sections, showed that XBP1s overexpression had no impact on normal myelination, with *XBP1^SC-OE^* being undistinguishable from WT nerves. Remarkably however, in the *S63del* model XBP1s overexpression appeared to ameliorate the hypomyelination defects with a clear trend towards reduced g-ratio values (**Fig. 6B-C**). A more detailed EM analysis confirmed improved myelination in *S63del/XBP1^SC-OE^*nerves when compared with *S63del*, with reduced g-ratio in axons with caliber between 2 and 5 μm (**Fig. 6D-E**). In addition, neurophysiological parameters were improved in adult *S63del* animals overexpressing XBP1s, with a significant increase in NCV and a strong trend towards reduction in F-wave latencies (**Fig. 6F-G**). Finally, these improvements were accompanied by a trend towards reduced levels of the negative regulator of myelination *Id2* (**Fig. 6H**) and a partial reduction in UPR markers belonging to the PERK pathway, as measured by WB for P-eIF2α and qRT-PCR for *Chop* and *Mthfd2* (**Fig. 6I-K)**, overall suggesting a readjustment in Schwann cell ER proteostasis.

**Figure 6.**
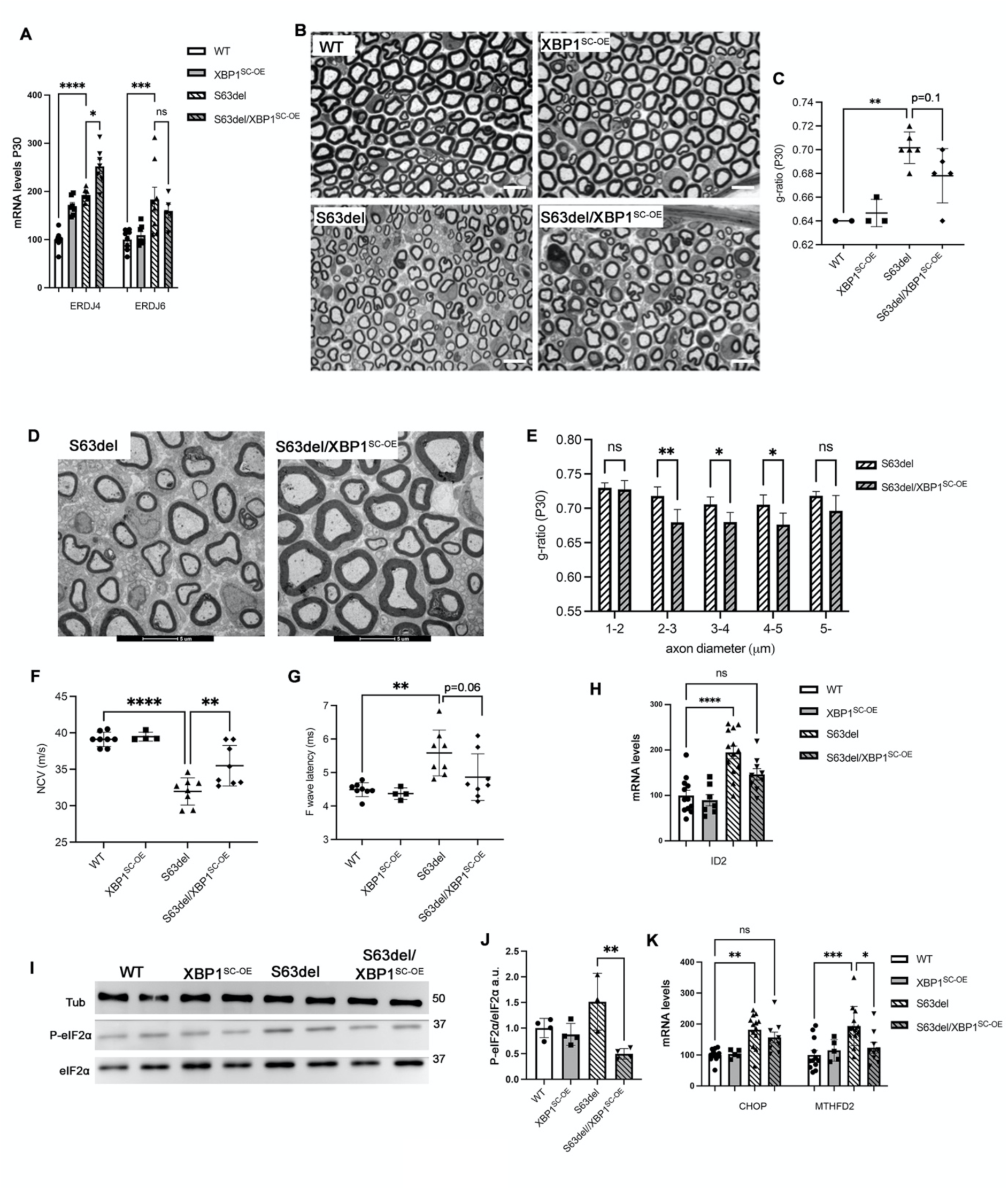
Schwann cell specific overexpression of spliced XBP1 ameliorates disease parameters in *S63del* mice. (**A**) qRT-PCR analysis on mRNAs extracted from P30 (upper graph) and P180 (lower graph) sciatic nerves for the XBP1s targets *Erdj4* and *Erdj6*. n = 6-8 RTs from independent nerves for each genotype. Values are expressed as fold change relative to WT (arbitrarily set to 100); error bars represent SEM. \*\*\*\**P* < 0.0001, * *P* <0.05 by one-way ANOVA followed by Tukey post hoc test. (**B**) Transverse semithin sections from P30 sciatic nerves. Scale bar 10μm. (**C**) G-ratio analysis performed on the P30 semithin sections. ** *P* <0.01 by one-way ANOVA (**D**) EM transverse section (scale bar 5μm) from P30 sciatic nerves and (**E**) relative g-ratio values plotted by axon diameter. Error bars represent SEM; ** *P* <0.01, * *P* <0.05 by unpaired Student’s *t*-test. n = 80-100 fibers per nerve were measured from 5 mice per genotype. (**F**) NCV and (**G**) F-wave latency measurements on 6-month-old mice; n = 4-8 mice per genotype. \*\*\*\**P* < 0.0001, ** *P* <0.01, by one-way ANOVA followed by Tukey post hoc test. (**H**) qRT-PCR analysis on mRNAs extracted from P180 sciatic nerves for the negative regulator of myelination *Id2*. *n* = 8-10 RTs from independent nerves for each genotype. Values are expressed as fold change relative to WT (arbitrarily set to 100); error bars represent SEM, *** *P* < 0.001 by one-way ANOVA followed by Tukey post hoc test. (**I**) Western blot analysis on P180 sciatic nerve extracts for P-eIF2α. Two representative samples out of four per genotype are shown; (**J**) quantification of P-eIF2α levels as arbitrary units (a.u.) relative to WT. (**K**) qRT-PCR analysis on mRNAs extracted from P180 sciatic nerves for *Chop* and *Mthfd2*. n = 5 RTs from independent nerves for each genotype. Values are expressed as fold change relative to WT (arbitrarily set to 100); error bars represent SEM, \*\*\*\**P* < 0.0001, *** *P* < 0.001, * *P* <0.05 by one-way ANOVA followed by Tukey post hoc test.

Importantly, comparable results were observed after crossing the more severe *R98C* model with *ROSA-XBP1s* mice. Adult *R98C/XBP1^SC-OE^* mice showed improved myelination, as measured by g-ratio analysis on both semithin and EM sections (**Fig. 7A-D**), with medium caliber axons in particular showing significantly reduced g-ratio values, and significantly increased NCV as compared to *R98C* mice (**Fig. 7E**). These ameliorations were associated to increased levels of P0 protein, pointing at improved myelination (**Fig. 7F-G),** to reduced levels of P-eIF2α protein (**Fig. 7F and H)**, and of the mRNA for *Chop* and *Mthfd2* (**Fig. 7I-H**), again suggesting reduced ER-stress levels. Taken together, these observations indicate that Schwann cell specific overexpression of XBP1s signaling has a therapeutic potential for multiple CMT1B neuropathies with activated UPR.

**Figure 7.**
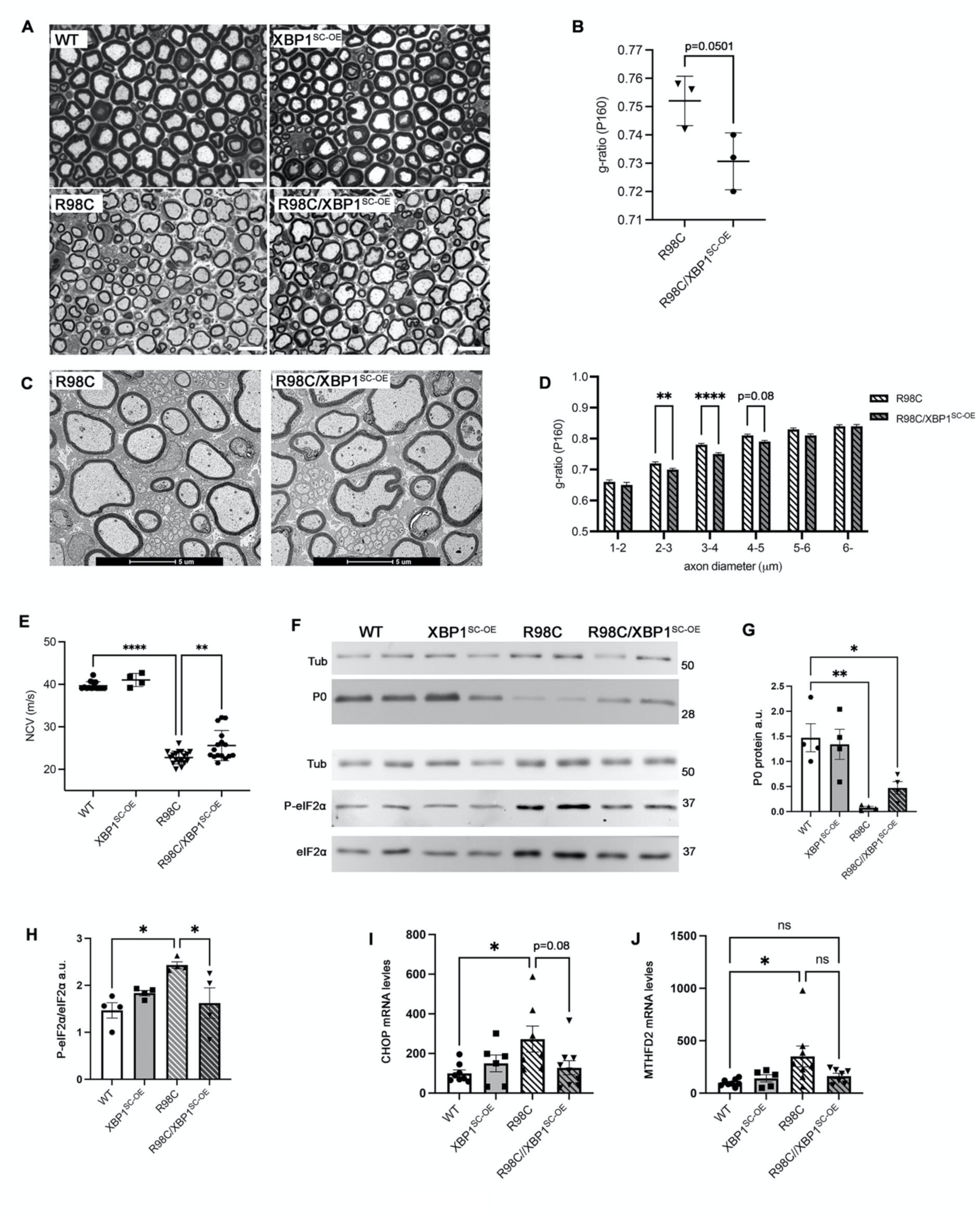
Schwann cell specific overexpression of spliced XBP1 ameliorates disease parameters in *R98C* mice. (**A**) Transverse semithin sections and (**B**) relative g-ratio analysis from adult (P160) sciatic nerves. (**C**) EM transverse sections (scale bar 5μm) and (**D**) g-ratio analysis from adult *R98C* and *R98C/XBP1^SC-OE^* sciatic nerves. n = 150 fibers per nerve were measured. \*\*\*\**P* < 0.0001, ** *P* < 0.01 by unpaired Student’s *t*-test. (**E**) Nerve conduction velocity studies. n = 4-12 mice per genotype. \*\*\*\**P* < 0.0001, *** *P* < 0.001 by one-way ANOVA followed by Tukey post hoc test. (**F**) Western blot analysis on sciatic nerve extracts for P0 and P-eIF2α; β-tubulin was used as loading control. Two representative samples out of four are shown for each genotype. (**G**) protein levels quantification for P0 and (**H**) P-eIF2α expressed as arbitrary units. Error bars represent SEM and ** *P* < 0.01, * *P* <0.05 by one-way ANOVA followed by Tukey post hoc test. (**I**) qRT-PCR analysis for *Chop* and (**J**) *Mthfd2* on mRNA extracts form P180 nerves. *n* = 8-12 RTs from independent nerves for each genotype. Values are expressed as fold change relative to WT (arbitrarily set to 100); error bars represent SEM, * *P* <0.05 by one-way ANOVA followed by Tukey post hoc test.

### Pharmacological IRE1/XBP1 activation ameliorates myelination in organotypic DRG cultures

Recently a series of IRE1α-XBP1 activators (IXA) have been identified.^50^ These compounds have been shown to selectively activate IRE1α RNAse activity and increase the amount of *Xbp1s* mRNA and downstream XBP1s target genes without activating the other UPR branches. Unfortunately, the current compounds are unable to pass the blood-brain and blood-nerve-barriers (BBB and BNB, respectively) (not shown). Thus, to test the potential of these compounds to treat CMT1B we turned once more to DRG explants. First, we confirmed that genetic overexpression of XBP1s would improve myelination also in *ex-vivo* organotypic cultures. Indeed, in DRGs from *S63del/XBP1^SC-OE^*mice, we observed an increased extent of myelination when compared to *S63del* (**Supplementary Figure 6A-B**), again with no alterations in WT myelination. Next, we evaluated the pharmacologic activation of the IRE1α-XBP1 pathway by testing compound IXA4^51^ on DRGs cultures from WT and *S63del* embryos (**Fig. 8A)**; qRT-PCR analysis confirmed increased expression levels for *Xbp1s* and its target *Erdj4* **(Fig. 8B)**. The treatment with the IXA4 compound was well tolerated, as shown by normal myelination in WT DRGs **(Fig. 8A and Supplementary Figure 6C-D).** Remarkably, in *S63del* DRG explants treatment with 1μM IXA4 improved myelination, as demonstrated by significantly increased number of MBP+ segments **(Fig. 8A and C)**, suggesting that pharmacologic activation of *Xbp1* splicing may represent a novel therapeutic avenue for CMT1B neuropathies.

**Figure 8.**
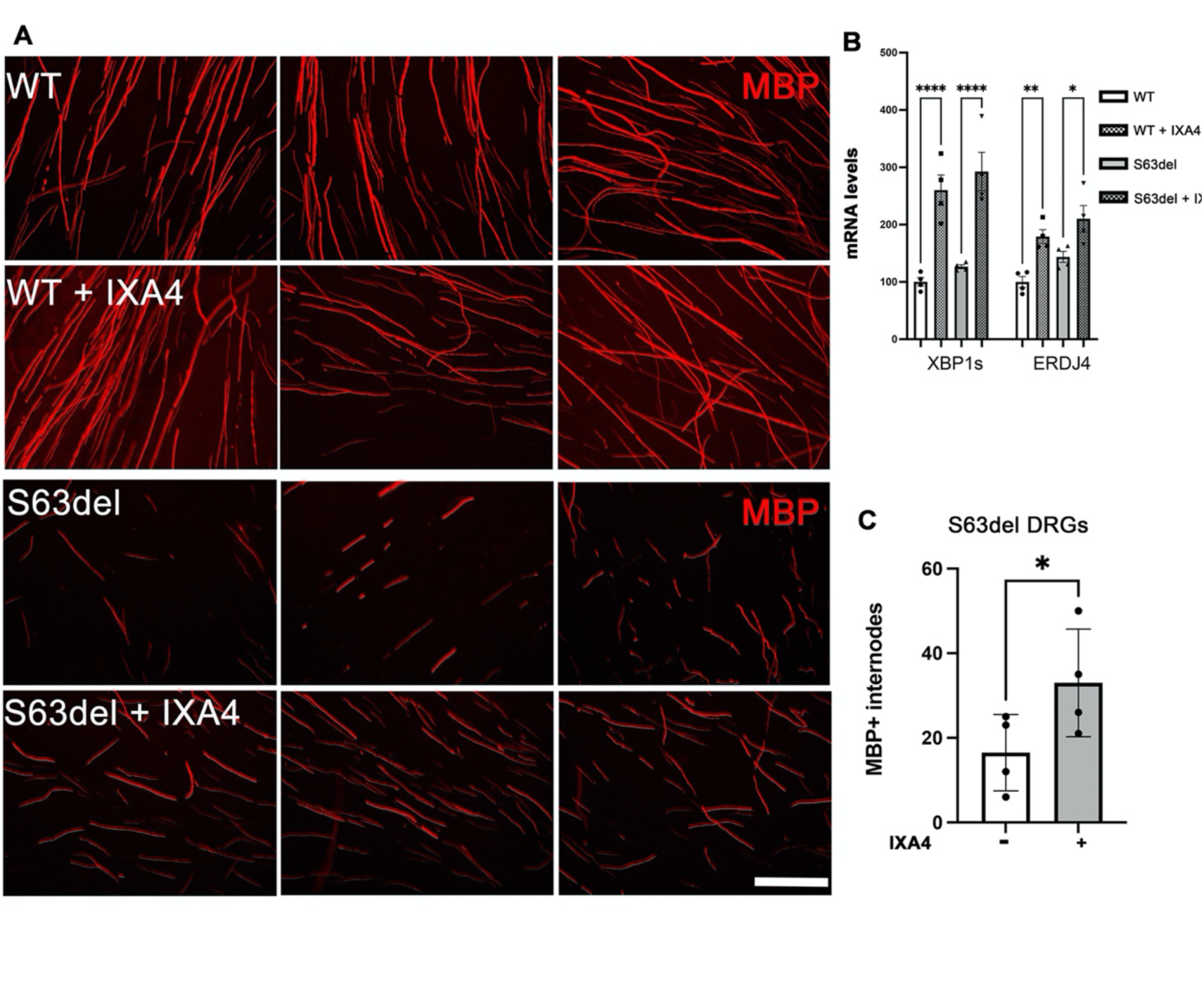
Pharmacologic activation of IRE1α mediated *Xbp1* splicing improves myelination in *S63del* dorsal root ganglia cultures. (**A**) Dorsal root ganglia were dissected from E13.5 WT and *S63del* embryos and myelination induced with 50μm ascorbic acid in the presence or absence of 1μm IXA4. Myelinating internodes were visualized with an antibody against myelin protein zero (MBP) (**B**) qRT-PCRs from mRNA extracted from WT or *S63del* cultures treated with vehicle or 1μm IXA4 for *Xbp1s* and its target *Erdj4*. *n* = 3-4 RT from independent experiments. Error bars represent SEM and \*\*\*\**P* < 0.0001, ** *P* < 0.01, * *P* <0.05 by one-way ANOVA followed by Tukey post hoc test. (**C**) Quantification of MBP+ internodes in S63del cultures after IXA4 treatment. * *P* <0.05 by Student *t*-test.

## Discussion

Defective protein folding and the activation of cellular stress pathways such as the UPR is a common mechanism in many myelin disorders, including CMT.^2,15^ We and others have previously shown the potential of modulating the PERK/P-eIF2α pathway for therapeutic intervention in various forms of CMT1,^17,18^ but the contribution of the other pathways of the UPR to neuropathy pathogenesis remained largely unexplored. Here we showed that XBP1 (the main effector of the IRE1α pathway) has a protective role in two different models of CMT1B, and that its genetic and pharmacologic activation represents a potential new therapeutic option for this currently uncurable disease.

### XBP1 is dispensable for Schwann cell development, myelination and remyelination after injury

Many studies have highlighted the importance of the IRE1α/XBP1 axis during development. Mice lacking *Xbp1* die around embryonic day 12-13.5 due to severe liver hypoplasia and fatal anemia.^22^ The generation of *Xbp1* conditional null mice has revealed the importance for this factor in the development of cell belonging to the immune system,^52^ and its crucial role in highly secretory cells such as B cells^23^ and Paneth cells^53^ as well as in lipogenesis^45^. During myelination Schwann cell need to synthesise large amounts of lipids and proteins that leads to a physiological activation of the UPR pathways.^54^ Yet, lack of *Xbp1* did not appear to impact on Schwann cell development and the myelination process. This is similar to what observed after Schwann cell specific ablation of *Perk*,^55^ impediment of eIF2α phosphorylation^43^ or ERAD impairment.^56^ These results suggest that Schwann cell are able to adapt to small fluctuations in cellular proteostasis, presumably by virtue of their remarkable plasticity^57^ and to efficient protein quality control systems.^2^

Recent work showed a specific activation of the IRE1α/XBP1 axis of the UPR in neuronal cell bodies and in Schwann cells of the distal segment of peripheral nerves after injury, and a decrease in locomotor recovery after nerve damage in mice lacking *Xbp1* in the nervous system.^38^ Overexpression of XBP1s, either in a mouse model where XBP1s is under the control of the prion promoter or through AAV mediated delivery of XBP1s into adult sensory neurons, accelerated axonal regeneration and locomotor recovery after nerve damage.^38^ Conversely, our results indicate that Schwann cell specific *Xbp1* depletion does not influence sciatic nerve remyelination after nerve crush, and thus suggest a neuronal cell autonomous role of XBP1 in nerve regeneration after damage.

### XBP1 is protective in different models of proteotoxic CMT1B

In stark contrast with its non-essential role in peripheral nerve myelination, maintenance and remyelination, we found that Schwann cell specific XBP1 expression is crucial to protect Schwann cells from extensive dysmyelination in two different models of CMT1B. In both *S63del* and *R98C* mice, ablation of *Xbp1* resulted in worsening of all the neuropathic features, from behaviour to neurophysiology to morphology. This is reminiscent of what observed in S63del mice carrying a non-phosphorylatable eIF2α allele (*S63del/eIF2α^SC-AA^* mice).^43^ Remarkably however, while the worsening of the phenotype in *S63del/eIF2α^SC-AA^* was mostly transient, so that adult *S63del/eIF2α^SC-AA^* were almost undistinguishable from *S63del* mice,^43^ *S63del/Xbp1^SC-KO^* and *R98C/Xbp1^SC-KO^* remain severely affected into adulthood, suggesting that a proper activation of the IRE1α/XBP1 pathway is even more crucial than the PERK/eIF2α pathway in the context of proteotoxic CMT1B.

XBP1s appears to exert its protective function mostly via up-regulation of genes involved in ER proteostasis pathways such as ERAD. This is particularly intriguing since we and others have recently shown that a functional ERAD is required to maintain myelin in adult glial cells,^56,58^ and that impairment of ERAD severely worsens myelin defects and neurophysiological parameters in CMT1B.^56^ Mutant misfolded P0 proteins are in fact degraded via the ERAD-proteasome pathway,^56,59^ and impairment of ERAD via Schwann cell specific ablation of Derlin-2 exacerbated the UPR and the overall phenotype of *S63del* mice.^56^ XBP1s appears to be upstream of this mechanism, and accordingly, ablation of *Xbp1* limits the activation of ERAD genes (including members of the Derlin family, such as Derlin-3), most likely resulting in an overwhelming accumulation of misfolded proteins in the ER of mutant Schwann cells. Indeed, in *S63del* nerves lacking *Xbp1* the transcriptional landscape is extensively altered: stress levels, as measured by targets of the PERK and ATF6 pathways are increased, but appear unable to compensate for the absence of XBP1s. This in turn leads the cells towards a maladaptive state, characterized by the activation of genes involved in cell death after prolonged UPR activation, such as *Trib3*, a target of the ATF4/CHOP axis,^60^ and of genes related to inflammatory processes, such as *Cxcl14* (see Fig. 4B). Of note, *Cxcl14* had been previously shown to be upregulated in CMT1A Schwann cells, and to affect Schwann cell differentiation and myelin gene expression.^61^ In this respect it is also noticeable that *S63del/Xbp1^SC-KO^* nerves show a significant increase in the expression of early Schwann cell transcription factors, such as Pou3f1/Oct6, Id2, Sox2 and c-Jun. The downregulation of Pou3f1 soon after birth is necessary for Schwann cell development,^62^ and its constitutive expression leads to hypomyelination.^63^ Similarly, hypomyelination was also a feature of mice overexpressing c-Jun^64^ that, like Id2 and Sox2, has been shown to work as a negative regulator of Schwann cell myelination.^40–42^ Increased levels of these factors have been previously detected in nerves of several neuropathic models, including *S63del*.^41^ Their extensive up-regulation in *S63del/Xbp1^SC-KO^* thus suggests that in conditions of unabated ER-stress Schwann cell differentiation is largely impeded. Intriguingly, a large increase of negative regulators of myelination and a block of Schwann cell differentiation was also detected in *S63del/eIF2α^SC-AA^* mice,^43^ where however lack of eIF2α phosphorylation did not result in hyperactivation of the other UPR pathways, suggesting different mechanisms in the control of Schwann cell proteostasis between the PERK and the IRE1α axis.

The ablation of *Xbp1* also led to an hyperactivation of IRE1α RNAse activity, which resulted in a supraphysiological splicing of the *Xbp1* mRNA lacking exon 2 and increased RIDD. This feedback loop had been previously observed in other models^32,45^ and indicates that XBP1 itself and the XBP1s-inducible genes have a strong suppressive function against IRE1α hyperactivation.^32,44^ This observation raised the question of whether IRE1α hyperactivity, through the induction of the JNK pathway or increased RIDD could contribute to the worsening of the CMT1B phenotype following *Xbp1* deletion. Our results however argue against this hypothesis; first, we did not detect any activation of the JNK/P-JNK pathway. Second, pharmacologic inhibition of IRE1α RNAse activity, and therefore of both *Xbp1* splicing and RIDD, had a similar effect as the *Xbp1* KO in myelinating *S63del* DRG cultures. These findings strongly suggest that the lack of the XBP1s is the major driver in the system, although only the full KO of IRE1α in CMT1B mice would unequivocally answer this question.

### Activation of XBP1s may represent a novel therapy for ER-stress related CMT

The modulation of protein quality control systems represents an appealing therapeutic candidate in CMT1 disorders characterized by the accumulation of misfolded protein.^65^ Curcumin, a low affinity SERCA inhibitor, has been shown to improve the trafficking of mutant proteins and reduce UPR markers in models of CMT1E and 1B.^66,67^ More recently, the direct modulation of UPR pathways was demonstrated to be effective in the amelioration of both CMT1B and CMT1A mice. Sephin1/IFB-088, a small molecule inhibitor of PPP1R15A/Gadd34, part of the complex that dephosphorylates eIF2α downstream of PERK, prolonged translational attenuation in stressed Schwann cells, allowing the cell to better dispose of mutant misfolded protein and to reset cellular proteostasis.^17,18^ Here we show that genetically potentiating XBP1s signalling specifically in Schwann cells was able to improve the phenotype of the *S63del* (CMT1B phenotype) and of the more severe *R98C* (DSS phenotype) mouse models. In both *S63del* and *R98C* mice we detected significant neurophysiological and morphological improvements. Due to the role of XBP1s in activating ER proteostasis genes, we speculate that this is likely due to improved handling of misfolded proteins within the ER. Although it is not possible to directly show reduction in mutant P0 accumulation in nerves *in vivo*, our data are supported by the reduction in overall stress levels which directly correlated to mutant protein expression.^8,16^ Consistently, molecules such as GlucNac, that promote general protein quality control system including autophagy and ERAD, were capable of improving myelination in DRG explants from CMT1B models,^56^ and more recently, elegant work has shown that raising cGMP and proteasomal activity ameliorated disease parameters in the *S63del* mouse.^59^ Of note, XBP1s overexpression improved *S63del* mice more than *R98C*. We posit two possible explanations for this: first, *S63del* mice present a milder phenotype to start with, which may be less difficult to improve as compared to the severe *R98C* pathology. Secondly, while *S63del* are transgenic mice that retain the two endogenous alleles of mouse *Mpz*, *R98C* are knock-in mice. Improved ERAD of the mutant protein would thus leave only one WT allele to produce the normal protein in *R98C* nerves, as compared to the two in *S63del*.

Importantly, in both *S63del* and *R98C* mice *Xbp1* mRNA is not fully spliced,^9,10^ and the remaining unspliced *Xbp1* may represent a therapeutic reserve. Accordingly, we also showed that the selective, pharmacologic activation of IRE1α/XBP1s largely increases *Xbp1* splicing and its target genes and can improve myelination in organotypic CMT1B DRG cultures. IXA4 has been shown to induce an adaptive remodeling in obese mice,^51^ affirming the potential of this strategy *in vivo*. The development of new IXAs capable of entering the nervous system is thus highly desirable as prospective novel treatment for several forms of demyelinating proteotoxic neuropathies.

## Acknowledgements

We thank Prof. Laurie H. Glimcher for the gift of the *Xbp1* floxed mice, and Dr. Cristina E. Barkauskas for the gift of the *ROSA26-hXBP1s* mice.

## Data availability

The data that support the findings of this study are available from the corresponding author, upon reasonable request.

## Supplementary material

Supplementary material is available online.

