## Supplemental Material (1 table, 6 Figures) for "Activation of XBP1s attenuates disease severity in models of proteotoxic Charcot-Marie-Tooth type 1B"

| | | |
|---|---|---|
| hXBP1F | human | AAACAGAGTAGCAGCTCAGACTGC |
| hXBP1R | human | TCCTTCTGGGTAGACCTCTGGGAG |
| mXBP1sF | mouse | GAGTCCGCAGCAGGTG |
| mXBP1sR | mouse | GTGTCAGAGTCCATGGGA |
| mXBP1totF | mouse | AAGAACACGCTTGGGAATGG |
| mXBP1totF | mouse | ACTCCCCTTGGCCTCCAC |
| CHOPF | mouse | GCGACAGAGCCAGAATAACAGC |
| CHOPR | mouse | TTCTGCTTTCAGGTGTGGTGGT |
| MTHFD2F | mouse | CATGGGGCGTGTGGGAGATAAT |
| MTHFD2R | mouse | CCGGGCCGTTCGTGAGC |
| BiPF | mouse | CTGAGGCGTATTTGGGAAAG |
| BiP78R | mouse | TCATGACATTCAGTCCAGCAA |
| GRP94F | mouse | CTCAGAAGACGCAGAAGACTCA |
| GRP94R | mouse | AAAACTTCACATTCCCTCTCCA |
| GADD34F | mouse | GAGGGACGCCCACAACTTC |
| GADD34R | mouse | TTACCAGAGACAGGGGTAGGT |
| ERDJ4F | mouse | CCATGAAGTACCACCCTGAC |
| ERDJ4R | mouse | CCTCTTTGTCCTTTGCCATTG |
| ERDJ6F | mouse | CGCCTTATCACAGTTTCACG |
| ERDJ6R | mouse | CTAAGAAGACGGTAGCTCTCC |
| BLOS1F | mouse | CAAGGAGCTGCAGGAGAAGA |
| BLOS1R | mouse | GCCTGGTTGAAGTTCTCCAC |
| Sel1LF | mouse | TGGGTTTTCTCTCTCTCCTCTG |
| Sel1LR | mouse | CCTTTGTTCCGGTTACTTCTTG |
| IRE1aF | mouse | CTGTGGTCAAGATGGACTGG |
| IRE1aR | mouse | GAAGCGGGAAGTGAAGTAGC |
| 36B4F | mouse | AGATTCGGGATATGCTGTTGG |
| 36B4R | mouse | AAAGCCTGGAAGAAGGAGGTC |
| XBP1 exon2/3 F | mouse | AAAACAGAGTAGCAGCGCAG |
| XBP1 exon2/3 R | mouse | AGCTGGAGTTTGTGGTTCTCT |
| Id2 F | mouse | TCCGGTGAGGTCCGTTAGG |
| Id2 R | mouse | CAGACTCATCGGGTCGTCC |

**Supplementary Table 1: qRT-PCR primer sequences**

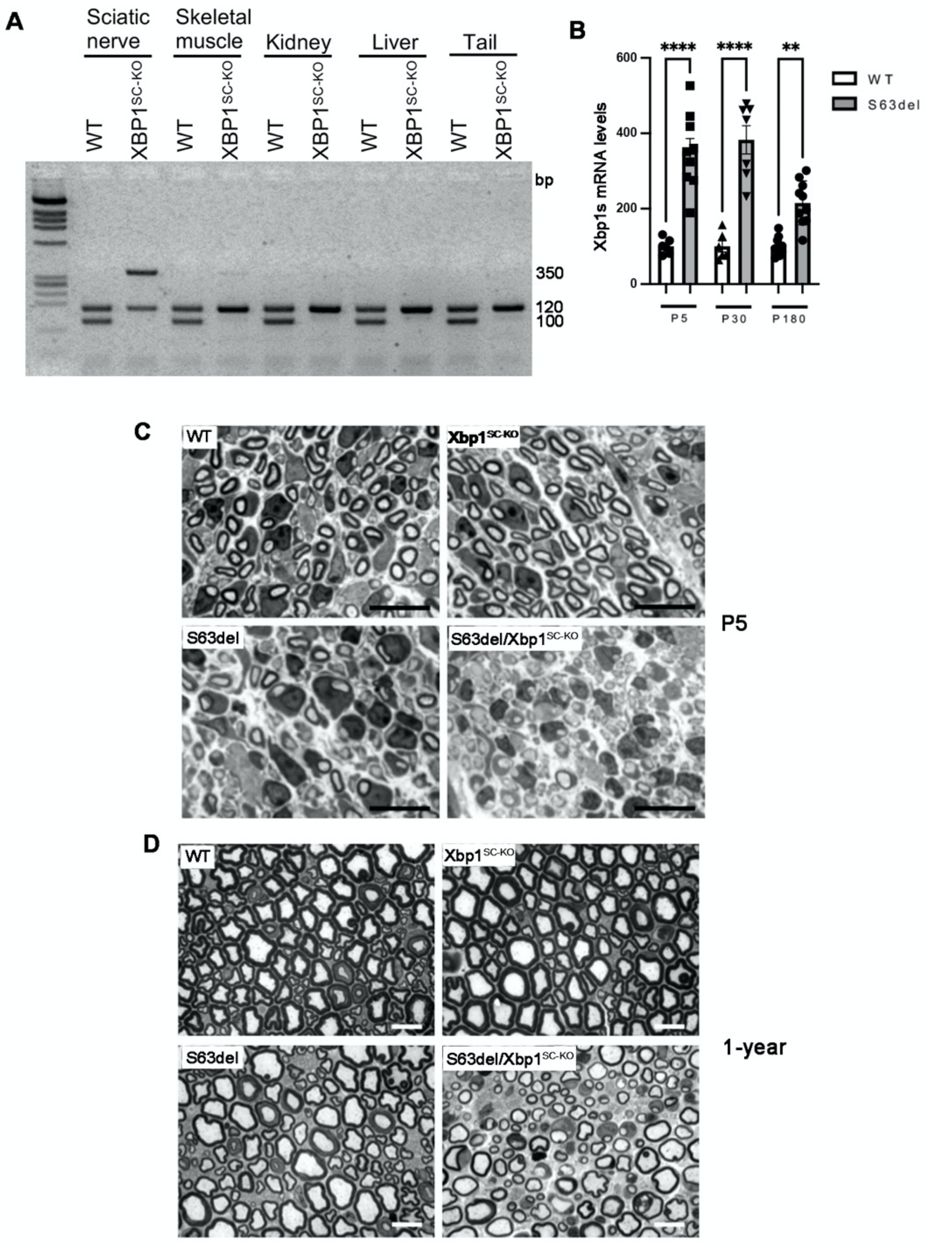


**Supplementary Figure 1**

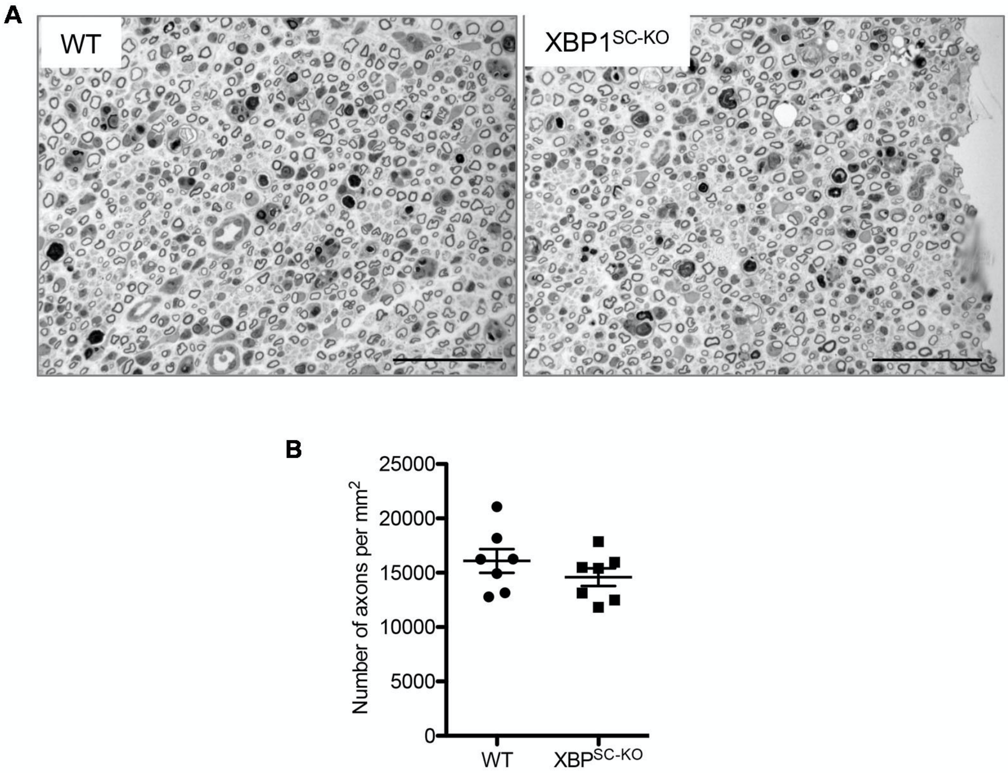


**Supplementary Figure 2**

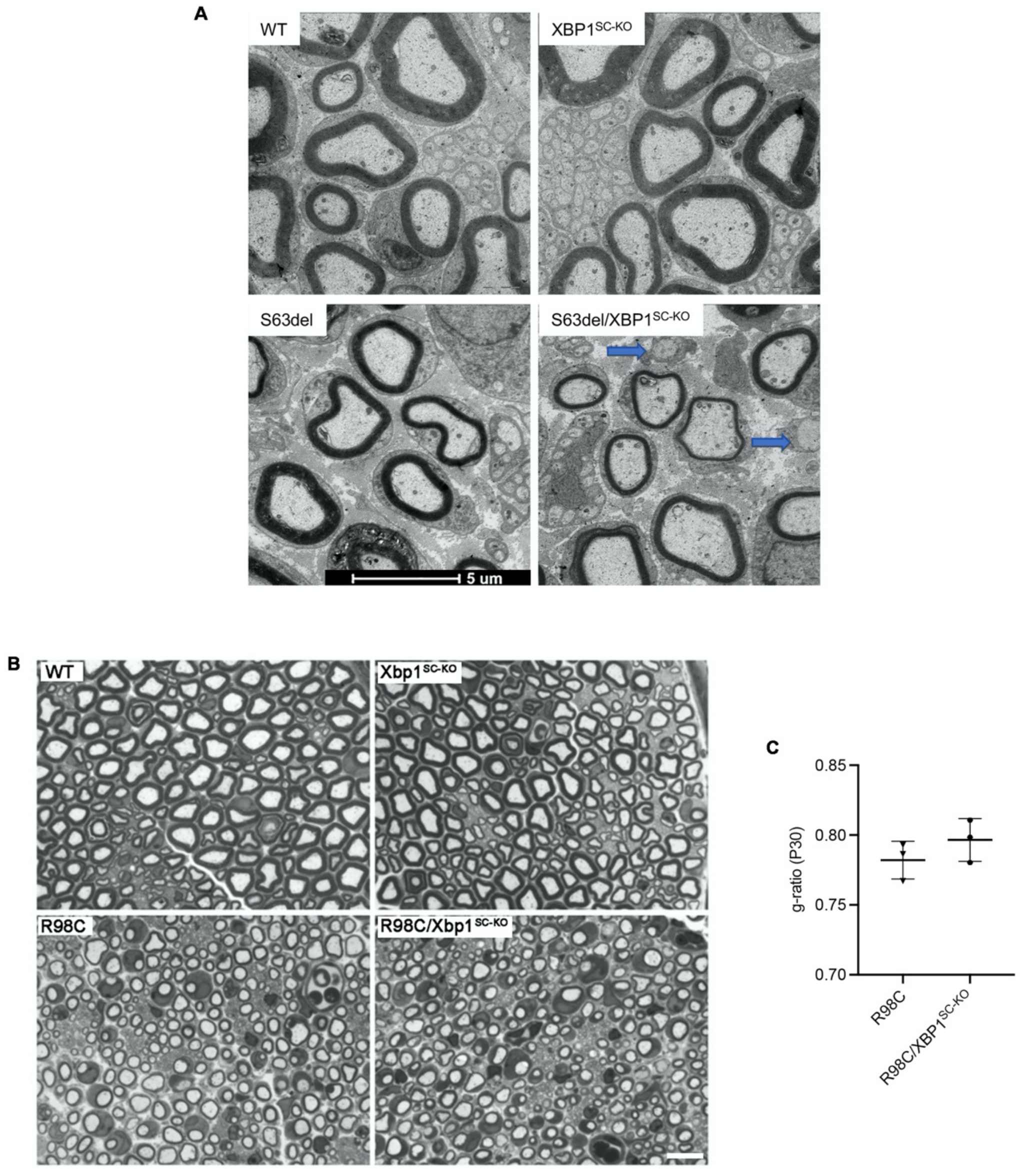


**Supplementary Figure 3**

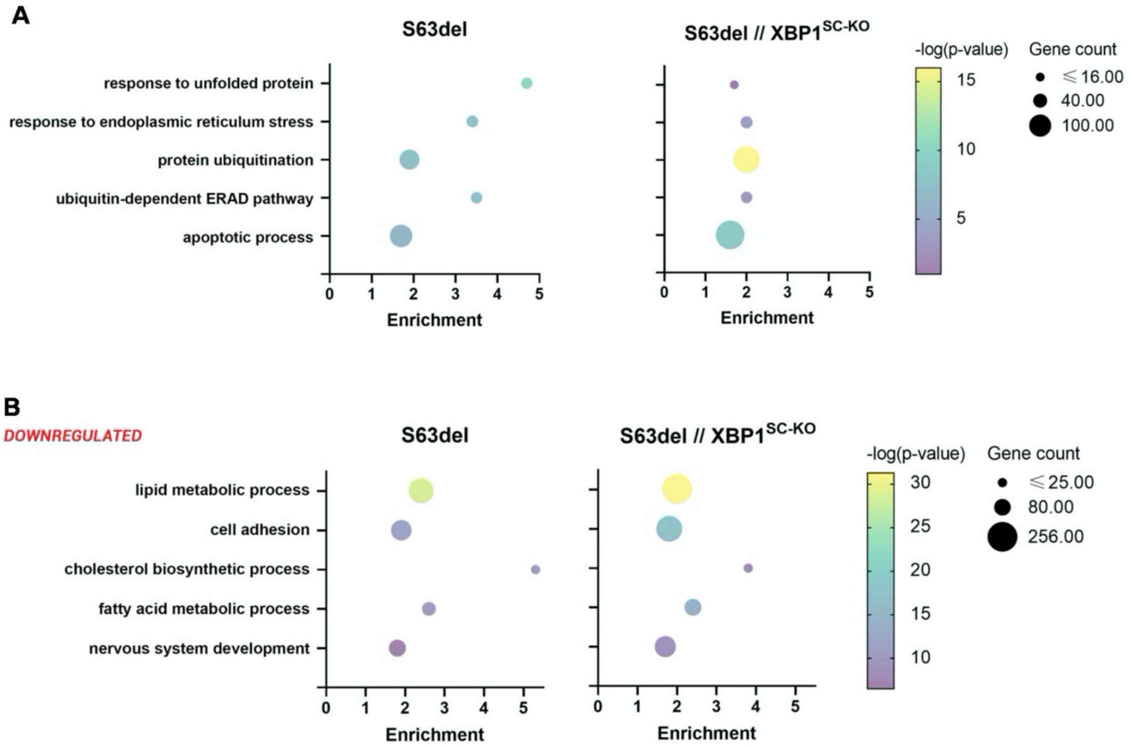


**Supplementary Figure 4**

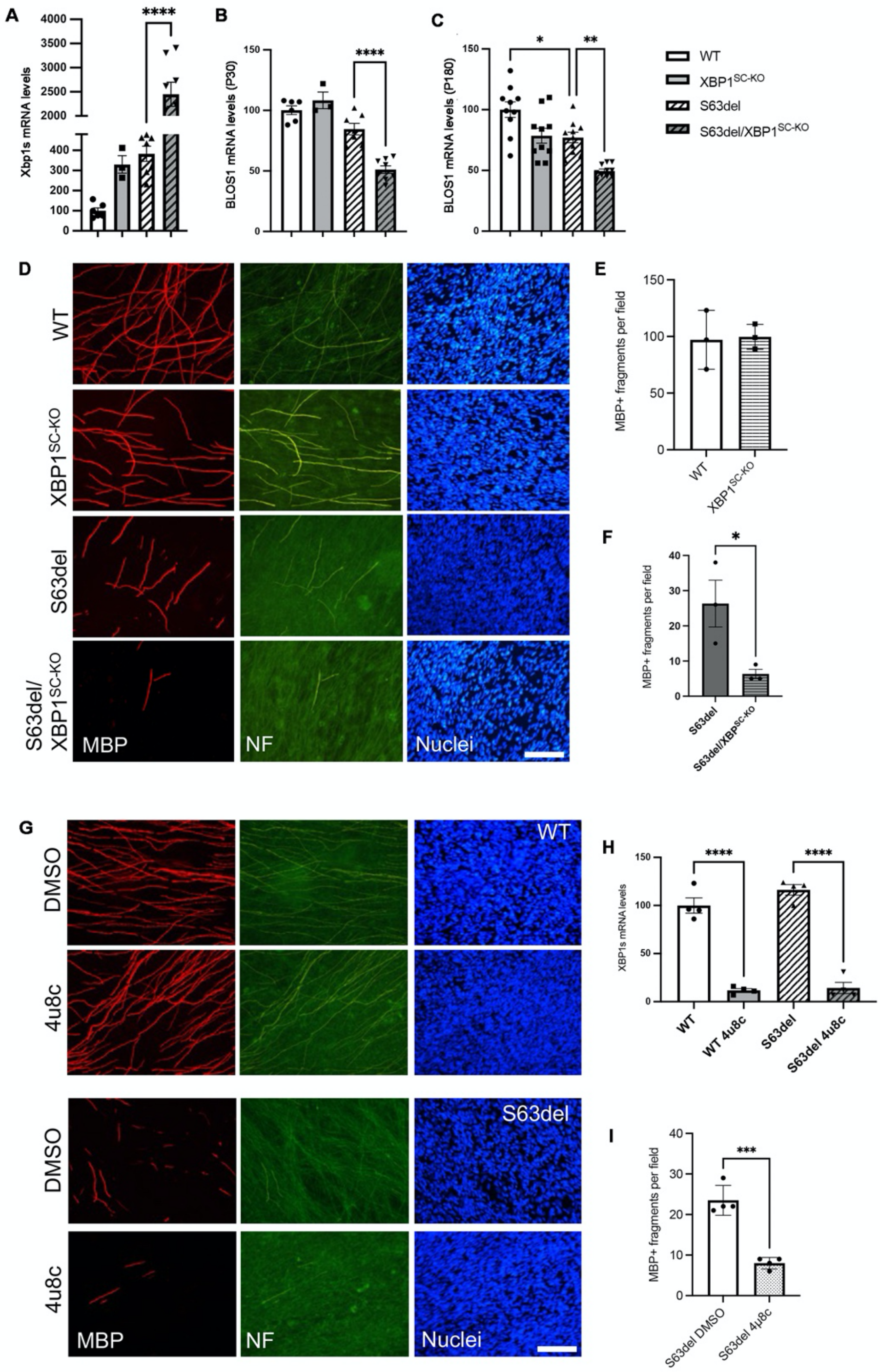


**Supplementary Figure 5**

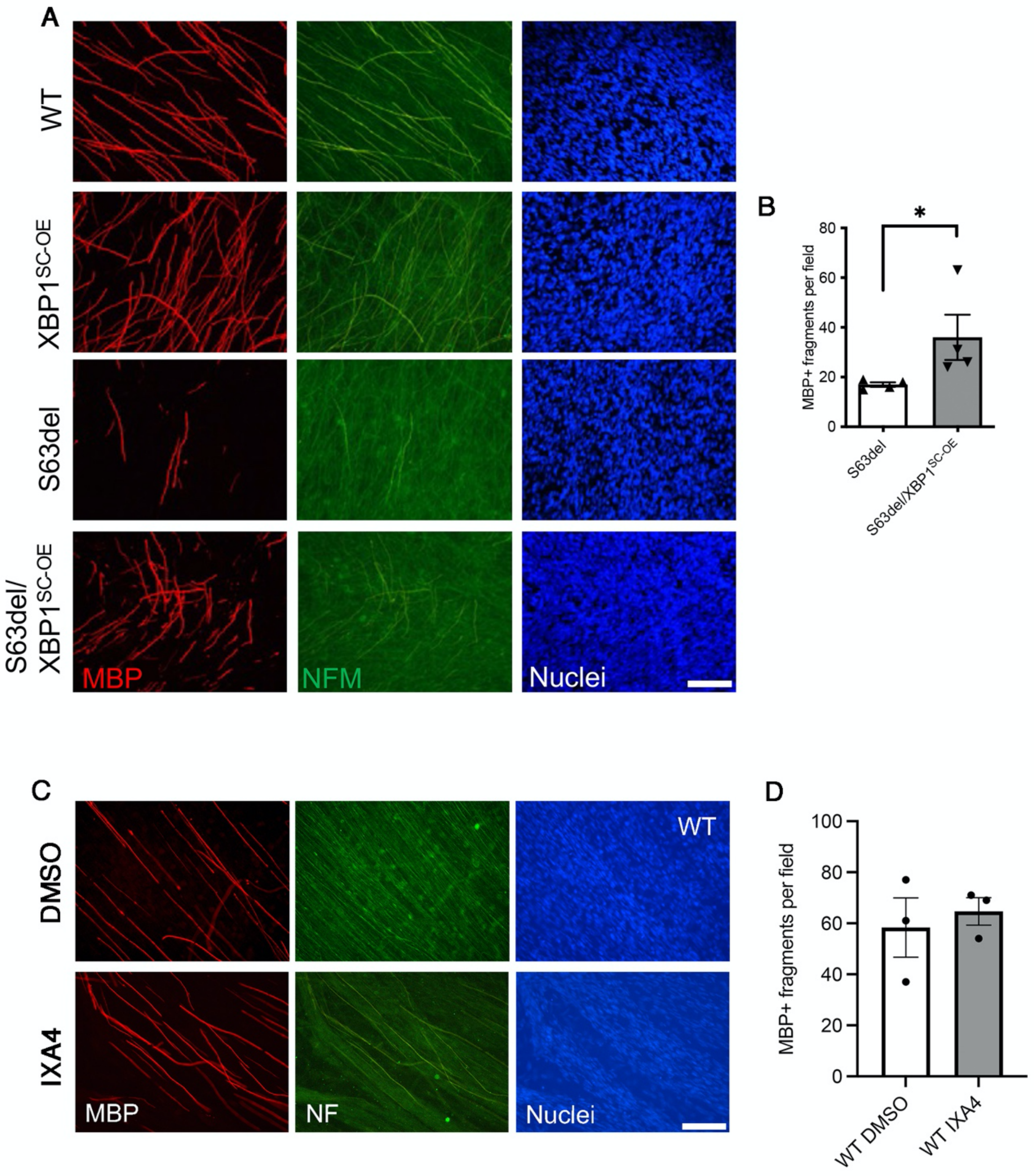


**Supplementary Figure 6**

### SUPPLEMENTARY FIGURE LEGENDS

**Supplementary Figure 1.** (**A**) PCR reaction on genomic DNA extracted from sciatic nerve and a selection of other tissues at P30. The 100 bp and 120 bp bands correspond to the WT and the floxed alleles respectively. A 350bp band corresponding to the recombined DNA is detected only in animals carrying the P0Cre. This band is detected only in sciatic nerves and not in other tissues. (**B**) qRT-PCR for spliced *Xbp1* on sciatic nerves from P5, P30 and P180 WT and *S63del* mice. Error bars represent SEM and ****$P < 0.0001$, **$P < 0.01$ by one-way ANOVA with Tukey post-hoc analysis, $n = 7$-9 RTs per genotype from independent nerves. (**C**) Transverse semithin sections of sciatic nerves from P5 and (**D**) 1 year old mice. Scale bar 10µm.

**Supplementary Figure 2.** (**A**) semithin section of sciatic nerves from WT and *Xbp1*$^{SC\text{-}KO}$ nerve 20-days after nerve crush. Scale bar 50µm. (**B**) quantification of the number of myelinated axons shows no differences amongst the two genotypes, $n = 7$ crushed nerves per genotype.

**Supplementary Figure 3.** (**A**) Electron microscopy on transverse sections of sciatic nerves from P30 mice. Note the hypomyelination in *S63del* nerves, which is further exacerbated in *S63del/Xbp1*$^{SC\text{-}KO}$ nerves were numerous amyelinated axons (blue arrows) could be detected. Scale bar 5µm (**B**) Semithin transverse sections from P30 WT, *Xbp1*$^{SC\text{-}KO}$, *R98C* and *R98C/Xbp1*$^{SC\text{-}KO}$ sciatic nerves. (**C**) g-ratio quantification for *R98C* and *R98C/Xbp1*$^{SC\text{-}KO}$ nerves, $n = 3$ nerves per genotype.

**Supplementary Figure 4.** (**A**) Gene Ontology enrichment for the processes related to UPR and ER-stress response in RNAseq data from *S63del* and *S63del/Xbp1*$^{SC\text{-}KO}$ nerves. (**B**) Gene ontology (GO) of biological processes downregulated in *S63del* (left) or *S63del/Xbp1*$^{SC\text{-}KO}$ (right) in comparison to WT.

**Supplementary Figure 5.** (**A**) qRT-PCR for spliced *Xbp1* on mRNA extracts from P30 sciatic nerves. Error bars represent SEM and ***$P < 0.001$ vs WT and $^{\#\#\#}P < 0.001$ vs *S63del* by one-way ANOVA followed by Tukey post hoc test; $n = 3$-7 RTs per genotype from independent nerves (**B**) qRT PCR for *Blos1* in P30 and (**C**) P180 mRNA extracts from sciatic nerves. Error bars represent SEM and ****$P < 0.0001$, ** $P < 0.01$ and * $P < 0.05$ by one-way ANOVA followed by Tukey post hoc test; $n = 3$-10 RTs per genotype from independent nerves. (**D**)

Dorsal root ganglia were dissected from E13.5 WT, *Xbp1*$^{SC\text{-}KO}$, *S63del* and *S63del/ Xbp1*$^{SC\text{-}KO}$ embryos and myelination induced with 50µm ascorbic acid. Myelinating internodes were visualized with an antibody against myelin protein zero (MBP). Antibodies for neurofilament (NF, green) and nuclei (DAPI, blue) were used to control for neuronal health and cell numbers (**E**) Quantification of MBP+ internodes in WT and *Xbp1*$^{SC\text{-}KO}$ cultures. (**F**) Quantification of MBP+ internodes in *S63del* and *S63del/ Xbp1*$^{SC\text{-}KO}$ cultures. Error bars represent SEM. $n = 3$ independent experiments; 4-8 DRGs per genotype were quantified per experiments. (**G**) DRG cultures from WT and *S63del* nerves treated with the IREα RNAse inhibitor 4u8c. (**H**) qRT-PCR for spliced *Xbp1* on mRNA extracts from the DRG cultures treated with 4u8c. Error bars represent SEM and ****$P < 0.0001$ by ANOVA followed by Tukey post hoc test, $n = 4$ DRGs per genotype. (**I**) Quantification of MBP+ internodes in *S63del* cultures after 4u8c treatment. Error bars represent SEM and ***$P < 0.001$ by Student's *t*-test. $n = 3$ independent experiments; 8-10 DRGs per condition were quantified per experiments.

**Supplementary Figure 6.** (**A**) Dorsal root ganglia were dissected from E13.5 WT, *XBP1*$^{SC\text{-}OE}$, *S63del* and *S63del/XBP1*$^{SC\text{-}OE}$ embryos and myelination induced with 50µm ascorbic acid. Myelinating internodes were visualized with an antibody against myelin protein zero (MBP). Antibodies for neurofilament (NF, green) and nuclei (DAPI, blue) were used to control for neuronal health and cell numbers. (**B**) Quantification of MBP+ internodes in S63del and *S63del/XBP1*$^{SC\text{-}OE}$ cultures. Error bars represent SEM and *$P < 0.05$ by Student's *t*-test. $n = 4$ independent experiments; 6-10 DRGs per condition were quantified per experiments. (**C**) DRGs from WT embryos were myelinated in the presence or absence of 1µm IXA4 for two weeks. Myelinating internodes were visualized with an antibody against myelin protein zero (MBP). Antibodies for neurofilament (NF, green) and nuclei (DAPI, blue) were used to control for neuronal health and cell numbers. (**D**) Quantification of MBP+ internodes from WT DRG cultures with or without IXA4. $n = 3$ independent experiments; 8-10 DRGs per genotype were quantified per experiments.
